# Bulk medial prefrontal cortex activity comports with a functional bipartition between salience detection and movement gain

**DOI:** 10.64898/2026.03.18.712525

**Authors:** Shikun Hou, Lindsey A Ramirez, Eric H Mitten, Jan Wiaderkiewicz, Manuela Narvaez Guzman, Heng Wang, Elizabeth J Glover

## Abstract

Reward prediction error (RPE) is a key function putatively realized by brain-wide neural states to drive learning and adaptive responding. The medial prefrontal cortex (mPFC) is among the many areas canonically implicated in RPE computation. Yet, whether mPFC RPE processing differs as a function of valence, value, and sign is not well-understood. Here, we performed *in vivo* mPFC calcium imaging in male and female rats engaged in an appetitive or aversive Pavlovian task designed to elicit RPE, along with continuous, unbiased behavioral monitoring. We found that short-latency bulk mPFC activity reports salience independent of valence, value, or RPE. Surprisingly, as subsequently validated with *in vivo* bidirectional optogenetics, we show that salience-null mPFC activity modulates generalized movement in a valence-and task-agnostic fashion. Together, these results challenge the pervasive notion of a unitary, region-wide representation of cognitive processes, including RPE signaling, by bulk mPFC activity, and instead, support a functional bipartition schema combining reactive detection of integrated salience and dynamic modulation of movement gain.

**Significance Statement:** By combining *in vivo* neural signal recording, bidirectional perturbation of activity, and continuous, unbiased, multidimensional behavioral segmentation during appetitive and aversive task performance, this study shows that region-wide bulk mPFC dynamics reflect a dual function of salience and movement encoding. These findings not only inform future work aiming to characterize mPFC dysfunction in disease, but importantly also highlight the crucial need for researchers to employ approaches that effectively distinguish cognition-like neural activity from movement-and sensory-induced dynamics.

## Introduction

Survival requires associative learning (e.g. stimulus-action-outcome), identification of salient predictors conveying divergent valence and value (e.g. conditioned stimuli), and adaptive behavioral re––sponding (e.g. approach, avoid) based on this information. As change is the law of nature, one’s internal model requires constant update to adapt accordingly, thus constituting additional learning. How the brain orchestrates this multi-step process has been an area of intense investigation for decades but is still not fully understood.

The reward prediction error (RPE) hypothesis^1, 2^ represents one of the first, and arguably the most influential, formalization of learning mechanics, which seeks not only to causally explain adaptive behavioral responding in terms of native neural dynamics, but also to model maladaptive behavioral deficits due to disrupted neural states. RPE, the perceived deviation of outcome value from its prediction – positive if better, and negative if worse – is hypothesized as an explicit driver of learning, whose implicit goal is error minimization^3^. Importantly, impaired RPE signaling has been implicated in a host of neuropsychiatric diseases including schizophrenia^4–7^ mood disorders^4, 5, 8, 9^ obsessive compulsive disorder^10^, and substance use disorders^3, 11–13^. Similar impairments have also been observed in neurological diseases including Parkison’s Disease^14^ and autism spectrum disorder^15^.

Since the original conception of the midbrain dopamine RPE hypothesis, mounting research has attributed wide-spread RPE encoding across many brain regions^16–18^. This includes the medial prefrontal cortex (mPFC)^11, 16, 18–30^ – a region characterized for its role in learning and adaptive behavioral responding^31, 32^ and whose function is significantly impaired in each of the aforementioned disease states^33–36^. However, the precise mechanisms by which mPFC assists in RPE computation as a function of valence (appetitive vs. aversive), value (good vs. bad), and sign (better vs. worse), are unclear, thereby precluding the ability of researchers to develop informed hypotheses regarding precise pathological dysfunction.

The present study sought to investigate the mPFC dynamics during appetitive or aversive learning and in response to expected and unexpected outcomes of either valence. Our results unexpectedly reveal that bulk mPFC Ca^2+^ activity encodes a combinatorial signal that indiscriminately reports the salience of environmental stimuli and movement independent of valence, value, and RPE. Using *in vivo* bidirectional optogenetics, our findings further demonstrate a causal link between salience-null mPFC activity and movement gain during, and beyond task-relevant performance.

## Materials and Methods

### Animals

All experiments used adult male and female Long-Evans rats (Envigo, Indianapolis, IN, USA) that were P60 upon arrival. The animal colony room was maintained on a 12-hour reverse light-dark cycle with lights on at 22:00. Upon arrival, rats were allowed to habituate to the colony room for at least one week before experiments began. Rats that underwent aversive Pavlovian conditioning had *ad libitum* access to food (Teklad 7912, Envigo) and water. Rats that underwent appetitive Pavlovian conditioning had *ad libitum* access to water and were food restricted to 95-100% free-feeding weight for at least five days prior to the start of behavioral testing. Food restriction was maintained at this level throughout the duration of the testing. All procedures were approved by the University of Illinois Chicago Institutional Animal Care and Use Committee and adhered to the NIH Guidelines for the Care and Use of Laboratory Animals.

### Stereotaxic surgery

Rats undergoing stereotaxic surgery were induced and maintained under a surgical plane of anesthesia using isoflurane (1-5%). Intracranial injections targeting the mPFC (AP: +3.2; ML: +1.4; DV: -3.5; 6° lateral) were made with back-filled custom glass pipettes connected to a nanojector (Drummond Scientific Company, Broomall, PA, USA) and administered at a rate of 10 nL/s. For *in vivo* photometry, rats of both sexes received a unilateral injection of 500 nL of AAV9-Syn-GCaMP7s-WPRE (Cat #104487; Addgene, Watertown, MA, USA) (**Figure 1a**). During the same surgery, a custom-made optical fiber (400 µm) was implanted just dorsal to the virus injection site and secured with dental cement. For optogenetics, male rats received bilateral injections of 500 nL of AAV8-Syn-ChR2(H134R)-GFP (Cat #58880; Addgene), whereas females were injected with AAV8-nEF-NpHR3.3-EYFP (Cat #137151; Addgene) (**Figure 7f**). A control group consisting of both sexes received bilateral injections of AAV2-hSyn-EGFP (Cat #50465; Addgene). During the same surgery, custom-made optical fibers (200 µm) were implanted bilaterally just dorsal to the virus injection sites and secured with dental cement. Rats were left undisturbed for at least three weeks before behavioral testing to allow for recovery and optimal viral expression.

**Figure 1:**
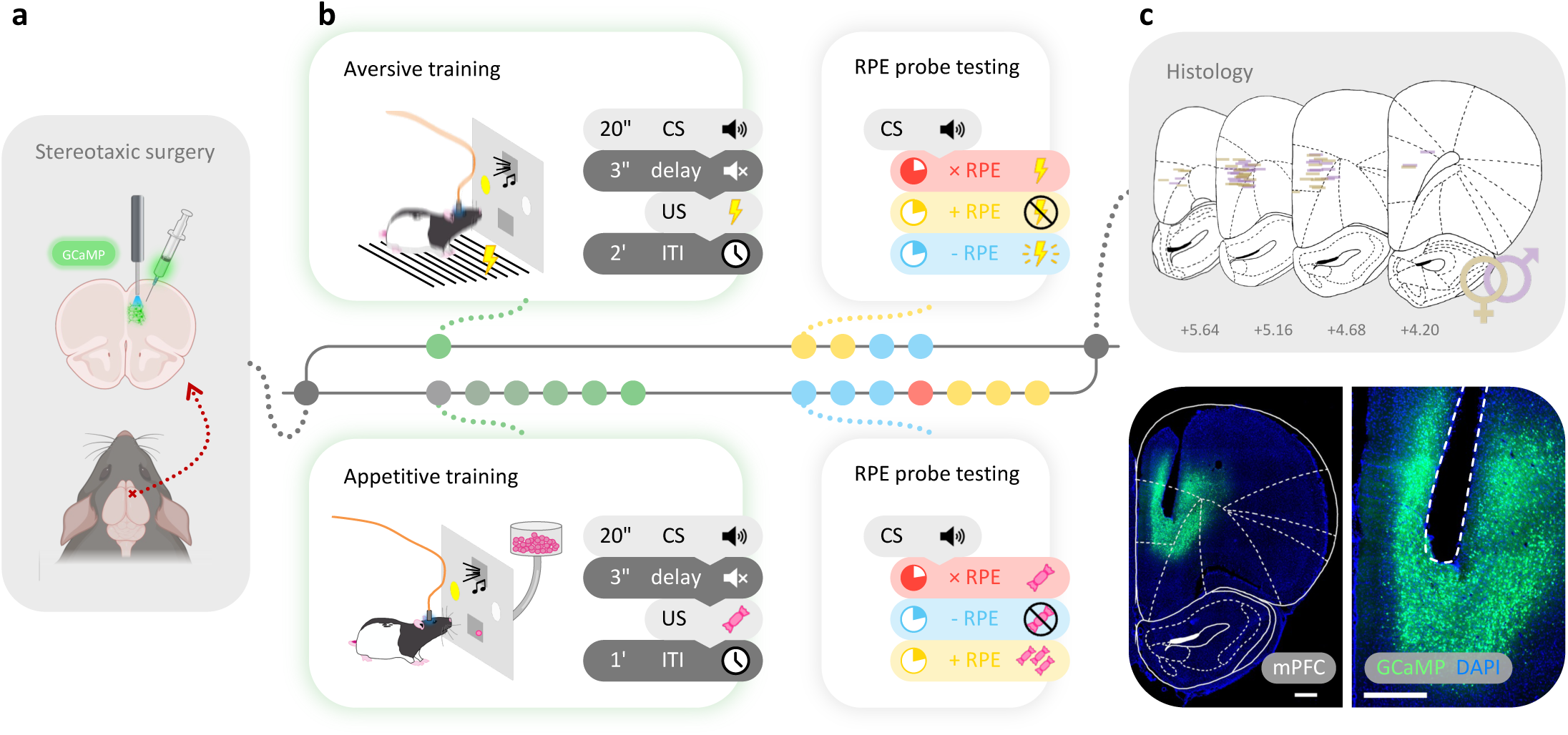
Using *in vivo* fiber photometry to record mPFC Ca^2+^ activity in response to unexpected, valenced outcomes. **a**, GCaMP7s was unilaterally expressed in the mPFC of male and female rats (n=66). **b**, Following recovery, rats underwent either aversive or appetitive Pavlovian conditioning to associate an audiovisual cue with the delivery of either a mild foot shock or a sucrose pellet. Following conditioning, rats underwent RPE probe test sessions during which outcomes that were better (+RPE) or worse (-RPE) than expected were delivered during 20% of the trials in a pseudo-random manner. Ca^2+^ fiber photometry and video recordings were collected across all stages of behavioral testing. **c**, Top: Optic fiber implant termination sites. Bottom: Representative image of GCaMP7s expression and implant track in the mPFC. scale bars = 500 μm.

### Behavioral testing

#### Apparatus and stimuli

Behavioral training and testing were conducted using standard operant boxes (Med Associates, St Albans, VT, USA) equipped with a red house light, cue lights, a tone generator, a sucrose pellet dispenser, and a grid floor capable of delivering electric foot shock. The food magazine where pellets were dispensed was equipped with an infrared beam to detect head entries. The house light was illuminated for the duration of all behavioral sessions.

#### Aversive Pavlovian conditioning

Following recovery from surgery, rats were habituated to daily handling and tethering. Rats assigned to the aversive task underwent one conditioning session to associate an audiovisual compound cue [conditioned stimulus (CS)] with the delivery of a mild foot shock [unconditioned stimulus (US)] (**Figure 1b, Top**) consisting of 10 trials, during which the presentation of a 20 s cue (2 or 8 kHz tone + light, delivered in either pips or constantly) was followed by the delivery of a mild foot shock (0.5 mA, 0.5 s). Cue presentation and shock delivery were separated by 3 s to allow for discrimination of Ca^2+^ signal associated with each stimulus. Trials were separated by a 120 s intertrial interval (ITI). Learning was indicated by an increase in conditioned freezing during the 20 s cue presentation and 3 s delay. After learning, rats underwent two positive reward prediction error (+RPE) probe sessions, each of which consisted of 30 CS-US pairings, where shock delivery was omitted pseudo-randomly during 20% of the trials. Rats then underwent two negative reward prediction error (-RPE) probe sessions where a foot shock stronger than that which was delivered during training (1.0 mA, 0.5 s) was delivered pseudo-randomly during 20% of the trials. Each aversive session was followed by a 30 min context-control session, where rats were allowed to roam freely inside the operant box (house light illuminated) without exposure to either CS or US, in order to reduce threat responding to the training/testing context.

#### Appetitive Pavlovian conditioning

Rats assigned to the appetitive task were trained during 6-9 conditioning sessions to associate the presentation of an audiovisual compound cue (CS) with delivery of a single sucrose pellet (US) (Bio-serv, Flemington, NJ, USA) (**Figure 1b, Bottom**). Each conditioning session consisted of 20 trials (60 s ITI) during which the cue was presented for 20 s followed by delivery of one sucrose pellet separated by a 3 s delay. Learning was indicated by an increase in anticipatory reward seeking during the 20 s cue presentation and 3 s delay. After training, rats were introduced to three -RPE probe sessions during which pellet delivery was pseudo-randomly omitted during 20% of the trials. Rats then underwent one day of remedial conditioning followed by three +RPE probe sessions during which 20% of trials resulted in pseudo-random delivery of three sucrose pellets.

### *In Vivo* Fiber Photometry

#### Ca^2+^ data acquisition

*In vivo* fiber photometry was used to collect Ca^2+^ signals and used as an indirect measure of neuronal activity in the rat mPFC (**Supplemental Movie 1**). Recordings were acquired during both conditioning and probe sessions. Ca^2+^ data were collected using either a Doric Fiber Photometry System (Sainte-Foy, QC, Canada) interfaced with a TDT RZ5P Fiber Photometry processor or a TDT RZ10X LUX Integrated Fiber Photometry Processor, both of which were controlled by the Synapse software (TDT, Alachua, Fl, USA). 465 nm light was used to excite Ca^2+^-dependent GCaMP7s fluorescence and 405 nm light was used as the isosbestic control for Ca^2+^-independent motion artifacts. TTL-pulses coincident with CS and US delivery were recorded to time-lock Ca^2+^ signal with task-relevant stimuli and behaviors.

#### Ca^2+^ data analysis

Daily photometry recordings were first processed in the open-source, Python-based, photometry toolbox, GuPPy^37^ to generate ΔF/F and z-scored peri-stimulus time histograms (PSTH) time-locked with stimuli and behaviors of interest. Briefly, 405 nm signal was first fitted to 465 nm signal using least-squares linear regression (**Supplemental Figure 1a-b**). The change in fluorescence (ΔF/F) was calculated by subtracting and dividing the fitted control from the Ca^2+^ signal (**Supplemental Figure 1c**) and z-scored to the mean and standard deviation of the entire trace. PSTHs were generated by isolating the z-scored ΔF/F around each event or behavior of interest and retimed so the time of 0 s aligned with the onset of a particular event or behavior. Baseline correction of PSTHs were used to control for trial-by-trial fluctuations in resting-state Ca^2+^ signal, by subtracting the mean z-scored ΔF/F during the 2 s baseline window prior to event onset from the PSTH of the entire trial.

Subsequent Ca^2+^ data processing, quantification, and analysis was performed with custom MATLAB (MathWorks Inc., Natick, MA, USA) scripts. Trial-level Ca^2+^ PSTHs were first down sampled to 60 Hz. Session-level PSTHs were averaged across multiple trial-level PSTHs of interest from the same session, and subject-level PSTHs were averaged across multiple session-level PSTHs of interest from the same subject. The slope of the subject-level PSTH was used as a summary measure to quantify the overall direction and degree of change of the Ca^2+^ activity of a predefined window, which was calculated by finding the coefficient of the least-squares linear fit and was used for statistical testing at the group level. Change from baseline was assessed by comparing the post-event slope with pre-event slope of the same duration. Group-level PSTHs shown in figures were averaged across multiple subject-level PSTHs of interest.

#### Signal-to-noise assessment

The quality of each photometry recording was quantified using a custom MATLAB script that measured signal-to-noise ratio (SNR). The power spectral density estimate (PSD) of a ΔF/F timeseries was obtained using the MATLAB “periodogram” function (**Supplemental Figure 1d**). The average power of the signal band of the PSD, defined as the frequency interval between 0 and 5 Hz, and the noise band, defined as the frequency interval above 5 Hz, was calculated using the MALTAB “bandpower” function. The SNR was the quotient of the signal band power and noise band power converted into the decibel (dB) scale. Recordings with a ΔF/F SNR lower than 29.6 dB (two standard deviations below the mean of 891 recordings) were considered low quality and therefore excluded from analysis (**Supplemental Figure 1e**). Subjects with more than 1/3 of recordings below threshold were excluded entirely.

#### Appetitive Ca^2+^ phenotyping

After all testing was completed, subjects in the appetitive task were classified *post hoc* as pellet-responsive if their average Ca^2+^ activity during the 2 s period following expected pellet delivery exhibited a positive slope, or non-responsive if they do not. To further support the validity of our Ca^2+^ phenotyping, we performed representational similarity analysis (RSA) inspired by previous reports^38, 39^ to confirm the binary grouping. The pairwise dissimilarity in average pellet related Ca^2+^ response was quantified by the correlation distance, which was taken as 1 - Pearson’s r, calculated using the MATLAB “corrcoef” function. The resulting cross-subject representational dissimilarity matrix (RDM) was visualized via multidimensional scaling (MDS) in a two-dimensional space using the MATLAB “mdscale” function (p = 2, Criterion = metricsstress). Stress was found to be 0.12 (< 0.2), suggesting a fair fit of data^39^.

### Optogenetics

#### Whole-cell patch-clamp slice electrophysiology

Rats were anesthetized with isoflurane and rapidly decapitated. Brains were removed and immediately placed in cold artificial cerebrospinal fluid (aCSF) cutting solution containing (mM): NaCl (125), KCl (2.5), NaH_2_PO_4_ (1.25), NaHCO_3_ (25), glucose (10), ascorbic acid (0.4), MgCl_2_ (4), and CaCl_2_ (1). Coronal slices containing the mPFC (300 μm) were made using a vibratome (Leica Biosystems, Wetzlar, Germany) and incubated at 34 °C in continuously oxygenated aCSF (95% oxygen/5% carbon dioxide) for at least 1 h prior to experiments. Following incubation, slices were transferred to a submerged recording chamber, held at 34 °C, and perfused at a rate of 2 mL/min with oxygenated recording aCSF solution containing (mM): NaCl (125), KCl (2.5), NaHCO_3_ (25), glucose (10), ascorbic acid (0.4), MgCl_2_ (1.3), and CaCl_2_ (2). The pH of all aCSF was adjusted to 7.2 – 7.4 and osmolarity was measured to be ∼300 mOsm.

Recordings were made with an Axon Multiclamp 700B amplifier, digitized at a sampling rate of 10 kHz with an Axon Digidata 1550A (Molecular Devices, San Jose, CA, USA) digitizer controlled by pClamp 11 software (Molecular Devices) running on Windows 10. Neurons were visualized using infrared interference-contrast microscopy (Olympus America, Center Valley, PA, USA), and recordings were obtained from GFP+ neurons in layer 5 of the mPFC ipsilateral to the injection site. Patch pipettes were constructed from thin-walled borosilicate capillary glass tubing (Warner Instruments, Holliston, MA, USA) and pulled with a P-97 pipette puller (Sutter Instruments, Novato, CA, USA) to a tip resistance of 3–5 MΩ. Membrane properties were assessed at break-in and at the end of each experiment. Any recordings for which the access resistance was >20 MΩ were discarded. Liquid junction potential was not assessed or corrected for in any experiment.

For current clamp experiments, patch pipettes were filled with a potassium gluconate intracellular solution containing (mM): K-gluconate (125), KCl (20) HEPES (10), ethylene glycol tetraacetic acid (EGTA, 1), magnesium adenosine triphosphate (Mg-ATP, 2), sodium guanosine triphosphate (Na-GTP, 0.3), phosphocreatine (10), and magnesium chloride (0.02), with pH adjusted to 7.3 and osmolarity measured at ∼285 mOsm. Baseline excitability was assessed by applying a series of 1 s current steps in 20 pA increments from −20 pA to 300 pA. After at least 1 minute of rest, the electrophysiological effects of channelrhodopsin (ChR2) and halorhodopsin (NpHR) activation on intrinsic excitability were validated using the same current step protocol in conjunction with light delivery occurring concomitant with current application. Light was generated by an LED equipped to the Olympus scope (470 nM: M470L5-C1, 530 nM: M530L4-C1; ThorLabs, Newton, NJ, USA), controlled by a T-cube LED driver (LEDD1B, ThorLabs), and delivered via a 40X objective to the slice bath. For ChR2, 470 nm light was delivered as pulse trains (10 Hz, 10 ms pulse width). For NpHR, 530 nM was delivered continuously. Light intensity for each wavelength was set at 10 mW.

#### In vivo optogenetic manipulation

For optogenetic stimulation, 450 nm light was delivered bilaterally (10 Hz, 10 ms pulse width) to ChR2-expressing rats. For optogenetic inhibition, 532 nm constant light was delivered bilaterally with laser offset ramped down linearly over 100 ms to prevent potential rebound excitation^40^. GFP-expressing controls received both 450 nm and 532 nm light delivery on separate sessions. Laser power was measured and adjusted before every session to target 8-10 mW optical power at the implant fiber tip.

After 6-8 weeks of post-surgery recovery and habituation, rats underwent aversive conditioning and +/-RPE probe testing sessions as described above (**Figure 7g**). After baseline testing, rats underwent two optogenetic manipulation sessions consisting of 30 cue-shock pairings during which light was delivered pseudo-randomly for 20 trials. Light onset occurred 2 s following cue onset and co-terminated with the 3 s delay preceding shock presentation. Light delivery also occurred during the ITI (≥ 30 s after shock presentation) in 10 trials. These sessions were followed by a context control session during which light was delivered for 21 s in a variable-interval (2.5 min) schedule for 10 trials in the absence of CS or US presentation (**Supplemental Movie 2**).

### Behavioral tracking and analysis

#### Video acquisition

Overhead videos of daily photometry sessions were recorded using a Pi USB Cam^41^ system equipped with infrared sensitivity and fisheye lens in conjunction with Ca^2+^ recordings interfaced with Synapse software (TDT, Alachua, Fl, USA; **Supplemental Movie 1**). Videos of daily optogenetics sessions were captured using the open-source video acquisition software OBS (Open Broadcaster Software) Studio at 640x480p in 20 FPS (**Supplemental Movie 2**). Video recordings from four subjects assigned to the appetitive task were captured using a commercial USB webcam and were subsequently excluded from all video-based behavioral analyses due to poor video quality, including low resolution and frame dropping as reported previously^41^.

#### Pose estimation

The open-source, deep-learning based, pose estimation platform SLEAP^42^ was used to track rat spatial location in acquired video recordings before subsequent behavioral quantification and classification (**Supplemental Figure 2a, d, Supplemental Movie 3, 4**). For photometry videos, a single-instance “unet” architecture was trained on 841 manually labeled frames from 28 videos containing male or female rats with diverse body sizes to track 8 key points (nose, head, right ear, left ear, shoulder, middle back, lower back, tail base). For optogenetics videos, a separate single-instance “unet” architecture was trained on 2,809 manually labeled frames from 70 videos to track the same set of key points.

#### Movement quantification

To quantify key-point tracking data from SLEAP and generate kinematics measures such as location and velocity, we adapted the open-source, MATLAB-based, behavioral analysis platform BehaviorDEPOT^43^. Briefly, instances of poor tracking results were first omitted, which for photometry videos, was achieved by removing frames with an “instance score” ≤ 2.5, and for optogenetics videos, by removing key points with “point score” ≤ 0.4 from a frame. Tracking data were then smoothed using a “LOWESS” smoothing method with a span of 5 frames, without interpolating missing points to prevent hallucination of key points that are deemed as occluded by SLEAP. Finally, the frame-by-frame linear velocity of each key point was calculated and stored for further analysis.

While event-specific behavioral summary measures were quantified and reported (e.g., mean velocity of shock response), behavioral PSTHs were constructed to examine the pattern and microstructure of behavioral responses to stimuli or behaviors of interest, similar to Ca^2+^ PSTHs. Frame-by-frame velocity time series aligned with stimulus or behavior timestamps were backward bin-averaged from 20 Hz to 4 Hz. The downsampled, trial-level, velocity PSTHs were averaged across trials, sessions, and subjects, to mirror the Ca^2+^ PSTH analysis.

#### Defensive behavior assessment and phenotyping

For photometry videos, passive (freezing) and active (darting) defensive behaviors were identified with the open-source, deep-learning based, behavioral classifier DeepEthogram^44^. Freezing was defined as the complete absence of movement except for respiration for a minimum of 0.5 s, and darting was characterized by sudden and rapid escape-like movement^45^ (**Figure 4a, Supplemental Figure 2a, Supplemental Movie 3**). We manually labelled 52 videos (∼70 hours) containing male or female rats with diverse body sizes using the DeepEthogram GUI for a total of 3 rounds of iterative training. The trained model was then used to generate inference on the rest of the dataset, and all predictions were manually examined and edited for errors. For optogenetics videos, defensive behaviors were identified with a custom MATLAB script using BehaviorDEPOT kinematic measures. First, occluded center and head velocities were interpolated using the MATLAB “fillmissing” function (‘movmedian’ method with a window of 80 frames). A freezing bout was identified if 1) both the head and center velocity were ≤ 0.5 cm/s, 2) the duration of the bout was ≥ 0.5 s, and 3) the inter-bout interval (IBI) was > 0.5 s. To identify darting, frame-by-frame center velocity acquired at 20 Hz was first bin-averaged to 5 Hz. Darts were identified using the MATLAB “findpeaks” on the down sampled velocity setting the 1) minimum peak darting velocity (‘MinPeakHeight’) to be 54 cm/s, 2) minimum inter-dart interval ‘(MinPeakDistance’) to be 0.4 s, and 3) minimum darting duration (‘MinPeakWidth’) to be 0.2 s. Identified darts were then classified as either *conditioned darting* if the dart occurred during cue presentation or the following delay period, or *shock response* if darting occurred during the 3 s after shock delivery. Timestamps of conditioned darting onset, freezing onset and offset were saved as .csv files for subsequent Ca^2+^ PSTH analysis. Freezing bouts whose offsets ended within ± 2 s of shock onset were excluded to prevent the pollution of shock-evoked Ca^2+^ activity in freezing-related Ca^2+^ analysis. Freezing probability PSTHs were generated by forward bin-averaging the freezing time series from 20 Hz to 4 Hz and analyzed similarly as the velocity PSTHs.

Upon the completion of all behavioral testing, rats in the aversive task were classified into the darter or freezer phenotypes using an estimated probability density function of the average count of conditioned darts per probe session (**Figure 4b**). A threshold of an average of 11 conditioned darts per probe session was determined by finding the first whole-number local minimum on a violin plot of all subjects’ performances using the MATLAB “kde” function. Rats with more darts on average per probe session than this threshold were classified as darters, and rats the fewer darts were classified as freezers.

#### Appetitive behavior assessment and phenotyping

Food magazine entry was measured by infrared beam break and used as an indicator of reward seeking. The first magazine entry during the 23 s period following cue onset was interpreted as the onset of anticipatory approach, whereas the first entry after pellet delivery was interpreted as the onset of reward retrieval and consumption. Entry timestamps were saved as .csv files for subsequent Ca^2+^ PSTH analysis. Continuous magazine entry probability was inferred from entry instances using a custom MATLAB script. Briefly, entry bouts were derived from entry instances where the start of a bout was identified as having a minimum inter-entry interval (IEI) of 2 s, and the end of a bout was defined as the last instance + mean IEI + 2 standard deviations. Assuming 100% entry probability within a bout, entry probability PSTHs in response to stimuli or behaviors of interest were forward bin-averaged to 4 Hz and analyzed similarly as all other behavioral and Ca^2+^ PSTHs.

Cue- and reward-oriented behavior was assessed using the locational proximity and orientation of the rat relative to tone speaker, cue light, and food magazine using a custom BehaviorDEPOT classifier (**Supplemental Figure 2d-f, Supplemental Movie 4**). Briefly, three regions of interest (ROI) around the food magazine, speaker, and cue light were drawn manually on each video. The rat was classified as interacting with an ROI if 1) the head was ≤ 5 cm from the ROI boundary, 2) the head angle was ≤ 45° from the ROI center, and 3) the interaction lasted ≥ 0.5 s. Food magazine ROI interaction was used as a proxy for reward-oriented behavior, while both speaker and light interaction were used as proxies for cue-oriented behavior. Behavioral probability PSTHs were analyzed similarly as above. A composite Pavlovian conditioned approach (PCA) index based on previously published work^46–48^ was calculated to quantify each rat’s bias toward either cue- or reward-oriented behavior during cue and reward period. Briefly, the PCA index score was the average of

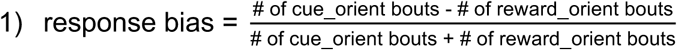

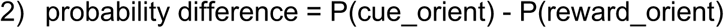

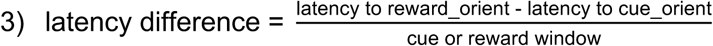

##### Histology

Rats were euthanized via transcardial perfusion. Photometry rats were perfused with phosphate buffered saline (PBS) containing 0.9 mM Ca^2+^ to enhance native GCaMP7s fluorescence followed by 4% paraformaldehyde (PFA). Optogenetics rats were perfused with PBS followed by 4% PFA. Brains were harvested and immersed overnight in 4% PFA followed by cryoprotection in 30% sucrose. Brains were then frozen on dry ice and stored at -80°C until ready for processing. Sections containing the mPFC were sliced coronally (40 µm) on a cryostat held at -20°C and then mounted onto frost plus slides and cover-slipped with Vectashield anti-fade mounting medium containing the nuclear stain, DAPI (Vector Laboratories, Burlingame, CA, USA). Viral expression and implant sites were confirmed visually via images acquired on a Zeiss AxioImager.M2 epifluorescent microscope (Zeiss, Oberkochen, Germany) (**Figure 1c, 7a,c**). Rats with misplaced implants were excluded from data analyses.

## Statistical analysis

Conventional statistical analyses were performed using GraphPad Prism 10 (GraphPad Software, Boston, MA, USA). For all analyses, significance level was set at P < 0.05 and data are presented as mean ± SEM unless stated otherwise. No statistically significant sex differences were observed with respect to either behavior or mPFC Ca^2+^ signal during training (**Supplemental Figure 3a-c**). Therefore, all analyses were collapsed across sex. Custom MATLAB scripts and R code used for behavior and Ca^2+^ analyses are available via our GitHub repository (<u>github.com/ejgloverlab</u>).

Cross-correlation between Ca^2+^ and velocity PSTHs was performed using a custom MATLAB script to quantify the relationship between Ca^2+^ and movement as a function of a time lag between the two time series. Ca^2+^ PSTHs were first downsampled to match the sampling rate (4 Hz) of velocity PSTHs. Due to the inherent time-dependent trend and therefore non-stationarity, PSTH time series were detrended via first differencing using the MATLAB “diff” function. Sample cross-correlation function (XCF) and corresponding lags between the differenced Ca^2+^ and velocity PSTH were calculated with the MATLAB “crosscorr” function specifying the maximum lag (‘NumLags’) to be ± 5 s and the number of standard errors in confidence bounds (‘NumSTD’) to be 1.96, which yields 95% confidence bounds assuming no correlation. A correlation is considered significant if it falls outside of the bounds. Cross-correlation was calculated for both the group mean PSTHs and subject mean PSTHs, and the subject sample XCFs were averaged across to generate the group average XCF. To statistically compare the relationships between Ca^2+^ and movement between groups, the maximum positive correlation with a lag within ± 3 s of each subject XCF was used for paired or unpaired t-test.

Because behavioral performance and Ca^2+^ signal were both highly dynamic across a second-to minute-long trial window and were measured according to a nested experimental design involving multiple trials, sessions, and subjects, functional linear mixed modeling (FLMM) was performed on select Ca^2+^ and behavioral PSTHs to uncover time-point-level and trial-level effects obscured by conventional summary measures of hierarchically averaged timeseries. The R package “fastFMM”^49^ was utilized in custom R scripts (version 4.4.2) to model the functional fixed-effects (regression coefficients denoted by ***β***(s)) of one or multiple covariates (e.g., trial type, phenotype, mean velocity, etc., denoted by **X***_i,j_*) on functional outcomes (e.g., Ca^2+^ or behavioral timeseries, denoted by **Y***_i,j_*(s), which is the outcome value for subject *i* at trial *j* at time *s*), while accounting for the subject-specific random intercept (denoted by ***γ****_0,i_*(s) for subject *i* at time *s*).

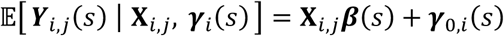

### *Velocity model (****Figure 7a***)

The effects of velocity on Ca^2+^ signal during aversive xRPE trials were modeled with:

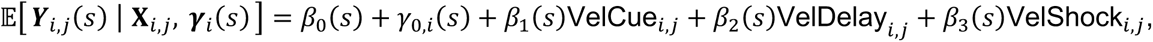

where the covariates were:

VelCue: mean velocity during cue (0-20 s) z-scored to all trials from all subjects,
VelDelay: z-scored mean velocity during delay (0-3 s) z-scored to all trials from all subjects,
VelShock: z-scored mean velocity during shock period (0-3 s) z-scored to all trials from all subjects.

Significant fixed-effects of each covariate was assessed for Ca^2+^ outcome of the corresponding trial period (e.g. association between VelCue and Ca^2+^ during cue).

### Optogenetics model (Supplemental Figure 9a)

The effects of optogenetic manipulation (main effects of and interaction between virus and laser) on behavioral performance during aversive optogenetic manipulation trials were modeled with:

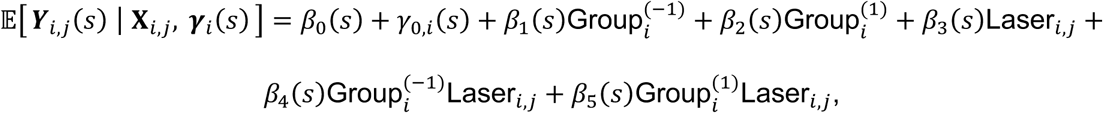

where the outcome **Y***_i,j_*(s) was either freezing probability or velocity, and the covariates were:
Group: factor variable, +1 to denote ChR2, -1 to denote NpHR, and 0 to denote GFP (reference level),
Laser: factor variable, +1 to denote laser ON trial, and 0 to denote laser OFF trial (reference level).

### Appetitive phenotype model of training (**Supplemental Figure 7a**)

The differences between appetitive phenotypes in Ca^2+^ and behavioral performance across training were modeled with:

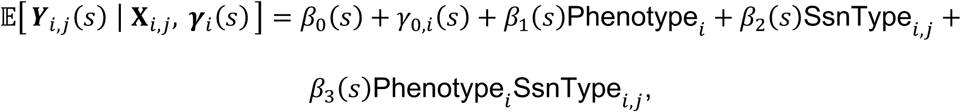

where the outcome **Y***_i,j_*(s) was Ca^2+^ or various behavioral timeseries, and the covariates were:

Phenotype: factor variable, 0 to denote pellet-responsive, and 1 to denote non-responsive (reference level),

SsnType: factor variable, 0 to denote first training session (reference level), and 1 to denote last training session.

### Appetitive phenotype model of -RPE (**Supplemental Figure 8a**)

The differences between appetitive phenotypes in Ca^2+^ and behavioral performance in response to reward omission were modeled with:

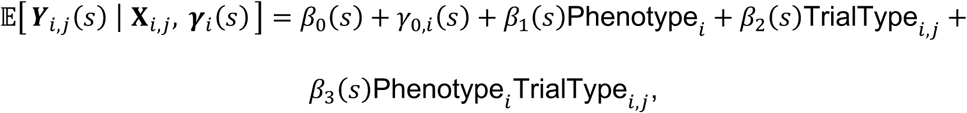

where the outcome **Y***_i,j_*(s) was Ca^2+^ or various behavioral timeseries, and the covariates were:

Phenotype: factor variable, 0 to denote pellet-responsive, and 1 to denote non-responsive (reference level),

TrialType: factor variable, 0 to denote -RPE reward omission trials (reference level), and -1 to denote the subsequent trials.

## Results

### mPFC Ca^2+^ activity reflects perceptual salience in a valence-non-specific manner

We sought to investigate whether and how region-wide mPFC encodes RPE by recording bulk pan-neuronal Ca^2+^ activity using fiber photometry while rats engaged in an aversive or appetitive Pavlovian task (**Figure 1**). As expected, conditioned freezing increased significantly across aversive conditioning trials during which an audiovisual cue was repeatedly paired with a mild foot shock (**Figure 2a,b**). Cue onset, offset, and shock presentation were each associated with a significant increase in mPFC Ca^2+^ activity from baseline throughout conditioning. Interestingly, we observed a significant attenuation of mPFC Ca^2+^ response to cue onset concomitant with potentiation of Ca^2+^ signal to cue offset across learning (**Figure 2a,c**). In addition, the shock-induced rise in mPFC Ca^2+^ was significantly more sustained after training (**Figure 2a,c**).

**Figure 2:**
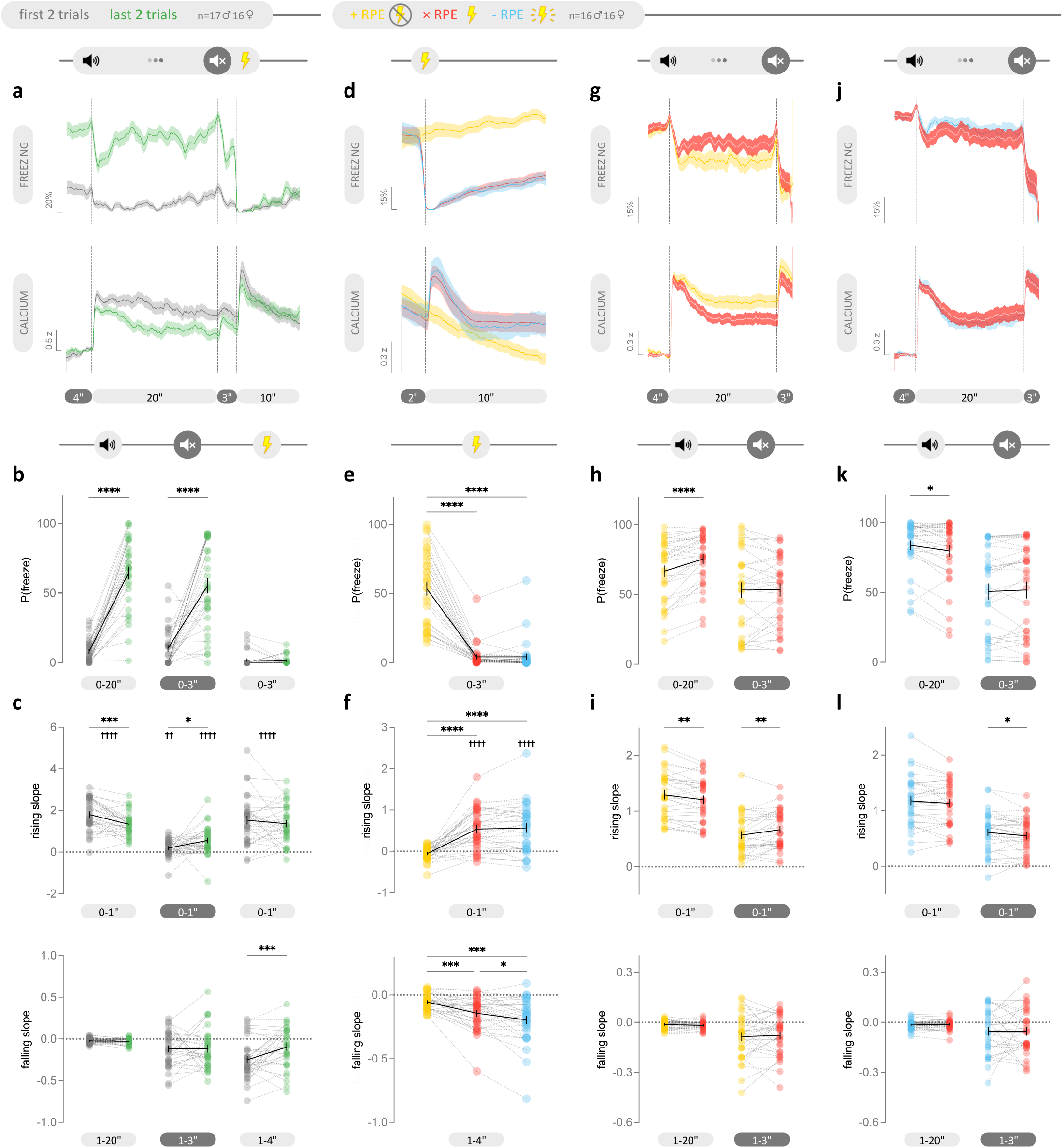
mPFC Ca^2+^ activity does not reflect aversive valence, value, or RPE. **a**, Mean freezing probability (top) and mPFC Ca^2+^ activity (bottom) in response to cue and shock presentation during the first (gray) and last two trials (green) of aversive conditioning. **b**, Rats froze significantly more during cue presentation (paired t test, P<0.0001) and following cue offset (paired t test, P<0.0001) after learning. **c**, Top: mPFC Ca^2+^ activity increased significantly from baseline in response to cue onset (paired t tests, first & last: P<0.0001), offset (paired t test, first: P=0.004, last: P<0.0001), and shock (paired t tests, first & last: P<0.0001). After learning, the increase in Ca^2+^ was attenuated in response to cue onset (paired t test, P=0.0003) but potentiated in response to cue offset (paired t test, P=0.0144). Bottom: The rate of decay in mPFC Ca^2+^ after shock presentation decreased significantly after learning (paired t test, P= 0.0007). **d**, Mean freezing probability (top) and mPFC Ca^2+^ activity (bottom) in response to expected mild shock (xRPE in red), unexpected shock omission (+RPE in yellow), and unexpected strong shock (-RPE in cyan). **e**, Rats froze significantly more during shock omission on +RPE trials compared to xRPE and -RPE trials (one-way RM ANOVA, P<0.0001; post hoc Tukey, P<0.0001). **f**, Top: mPFC Ca^2+^ activity significantly increased in response to mild and strong shock delivery compared to baseline (paired t test, P<0.0001) and shock omission (one-way RM ANOVA, P<0.0001; post hoc Tukey, P<0.0001). Bottom: Ca^2+^ decay in response to shock differed across RPE outcomes (one-way RM ANOVA, P<0.0001). **g**, **j**, Mean freezing probability (top) and mPFC Ca^2+^ activity (bottom) in response to cue for +RPE (**g**, yellow) or -RPE (**j**, cyan) trials and the trials (red) immediately following. **h**, Rats increased freezing during cue presentation on trials following a +RPE trial (paired t test, P<0.0001) **i**, Top: This was associated with attenuated Ca^2+^ response to cue onset (paired t test, P=0.0028) but potentiated Ca^2+^ response to cue offset (paired t test, P=0.0057). Bottom: The rate of Ca^2+^ decay during cue presentation was not different between +RPE and xRPE trials. **k**, In contrast, freezing was significantly decreased on trials that followed a -RPE trial (paired t test, P= 0.0143). **l**, Top: -RPE outcomes attenuated Ca^2+^ response to cue offset (paired t test, P=0.0324). Bottom: Similar to +RPE, Ca^2+^ decay was not different between -RPE and xRPE trials. PSTHs are expressed as group mean ± shaded SEM. Before-after plots show both group mean ± SEM (black) and individual data as colored dots with lines (dotted lines=males; solid lines=females). \**P* < 0.05, \*\**P* < 0.01, \*\*\**P* < 0.001, \*\*\*\**P* < 0.0001; ††*P* < 0.01, ††††*P* < 0.0001 from baseline.

To examine changes in mPFC activity in response to unexpected aversive outcomes, rats underwent additional testing during which 20% of trials were associated with an unexpectedly better (no shock; +RPE) or worse (stronger shock; -RPE) outcome (**Figure 1b** **top**). As expected, unlike -RPE and no RPE (xRPE) trials, rats maintained high levels of freezing during shock omission on +RPE trials (**Figure 2d,e**). mPFC Ca^2+^ activity significantly increased in response to both mild and strong shock, but the magnitude of increase did not scale with the magnitude of outcome value or RPE (**Figure 2d,f**). In contrast, mPFC Ca^2+^ did not change from baseline during +RPE trials (**Figure 2d,f**). The differences in Ca^2+^ kinetics associated with RPE outcomes were also reflected in the rate of decay (**Figure 2d,f**). Analysis of trial-by-trial changes in behavior and mPFC Ca^2+^ revealed potentiation of threat responding on trials immediately following +RPE trials (**Figure 2g,h**) and attenuation after -RPE trials (**Figure 2j,k**), suggestive of bidirectional changes in perceived threat in accordance with the sign of the PE outcomes. In a manner consistent with changes in mPFC Ca^2+^ observed as perceived threat was updated over the course of training, mPFC Ca^2+^ response to cue onset was attenuated following +RPE trials (**Figure 2g,i**), but potentiated during cue offset after +RPE trials (**Figure 2g,i**). In contrast, mPFC Ca^2+^ response to cue offset was attenuated after -RPE trials (**Figure 2j,l**). Decay kinetics during cue presentation were not dynamically responsive to trial-by-trial changes in aversive outcomes (**Figure 2i,l**). In summary, we did not observe a shift in mPFC response bias from US to CS across learning, nor a bidirectional modulation of activity by RPE outcomes of opposing signs, as would be predicted by RPE encoding. Moreover, +RPE outcomes resulted in similar directional changes in cue response as training, whereas -RPE outcomes resulted in the opposite, again incongruent with RPE encoding. Instead, the indiscriminate response to environmental stimuli of varying value, valence, and predictability seems to reflect perceptual salience encoding.

To examine whether mPFC activity differs in response to rewarding outcomes, mPFC Ca2+ dynamics were captured in a separate group of rats trained to associate cue presentation with delivery of a sucrose pellet. As expected, rats exhibited a significant increase in conditioned reward seeking over the course of training with seeking increasing in proximity to reward delivery (**Figure 3a,b**). Similar to our observations in the aversive task, mPFC Ca^2+^ activity significantly increased in response to cue presentation and reward delivery, with responses during cue onset and offset decreasing and increasing, respectively, across conditioning sessions (**Figure 3a,c**). mPFC Ca^2+^ response to appetitive cue offset also showed faster decay across conditioning sessions (**Figure 3a,c**).

**Figure 3:**
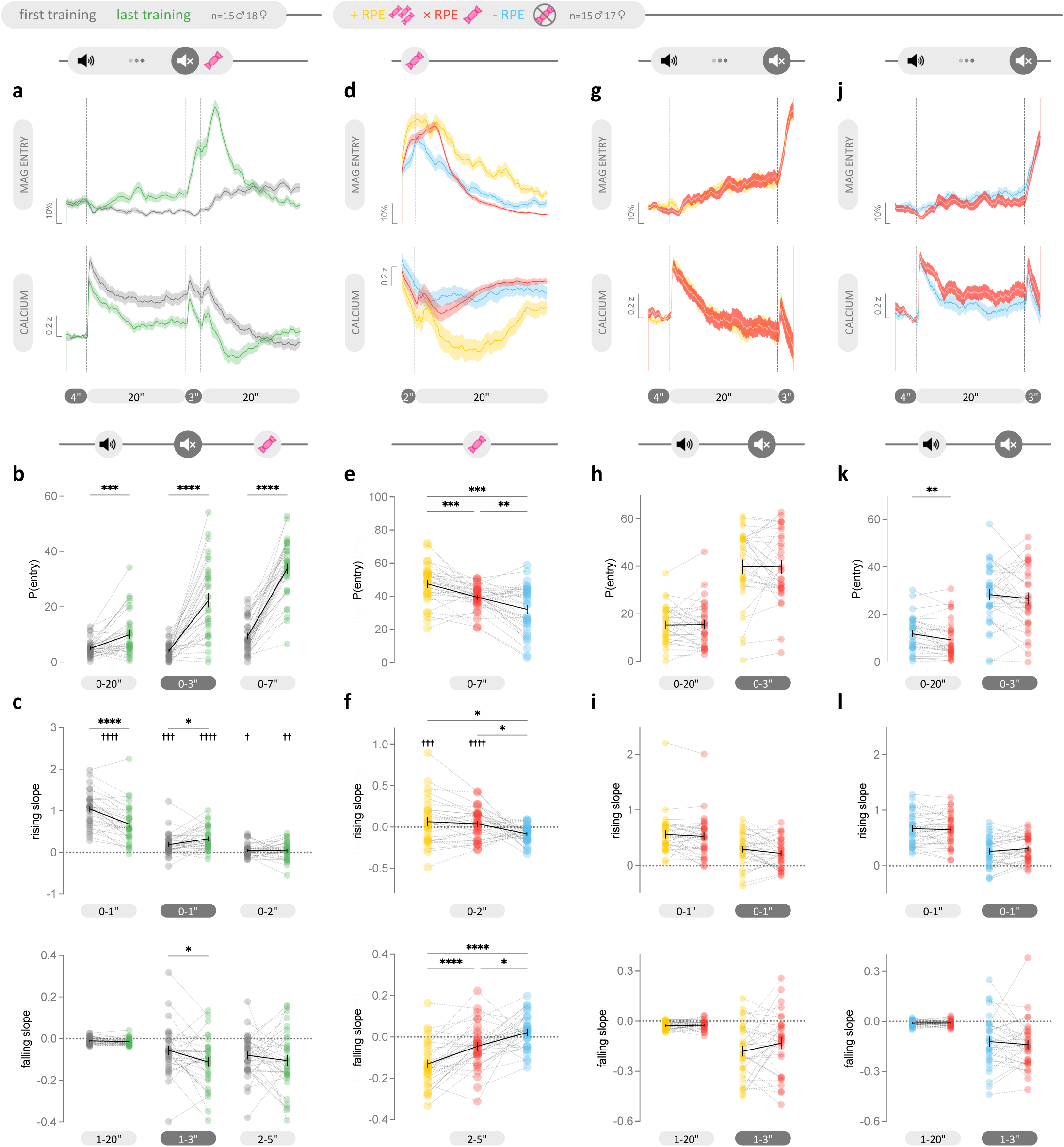
mPFC Ca^2+^ activity does not reflect appetitive valence, value, or RPE. **a**, Mean magazine entry probability (top) and mPFC Ca^2+^ activity (bottom) in response to appetitive cue and pellet delivery during the first (gray) and last conditioning session (green). **b**, Rats exhibited higher probability for magazine entry during cue presentation (paired t test, P<0.0002), the delay between cue and reward (paired t test, P<0.0001), and reward delivery (paired t test, P<0.0001) after learning. **c**, Top: mPFC Ca^2+^ activity significantly increased from baseline in response to cue onset (paired t test, P<0.0001), offset (paired t test, first: P=0.0003, last: P<0.0001), and pellet delivery (paired t test, first: P=0.0120, last: P=0.0021). After learning, the increase in mPFC Ca^2+^ was attenuated in response to cue onset (paired t test, P<0.0001) but potentiated in response to cue offset (paired t test, P=0.0107). Bottom: The rate of decay in Ca^2+^ during the delay period increased significantly after learning (paired t test, P=0.0219). **d**, Mean magazine entry probability (top) and mPFC Ca^2+^ activity (bottom) in response to appetitive cue and pellet delivery in response to expected reward (xRPE in red), unexpected larger reward (+RPE in yellow), and unexpected reward omission (-RPE in cyan). **e**, Rats engaged in more magazine entries on +RPE trials (mixed-effects analysis, P<0.0001; post hoc Tukey, P=0.0004) and fewer entries during -RPE trials (post hoc Tukey, P= 0.0048) relative to xRPE trials. **f**, Top: mPFC Ca^2+^ activity significantly increased in response to small and large rewards compared to baseline (paired t test, small: P<0.0001, large: P=0.0008) and to reward omission (mixed-effects analysis, P=0.0113; post hoc Tukey, xRPE: P=0.022, +RPE: P=0.0450). Bottom: Greater decay in Ca^2+^ activity was observed during large reward delivery (mixed-effects analysis, P<0.0001; post hoc Tukey, P<0.0001) and less during reward omission (post hoc Tukey, P=0.0213) compared to xRPE trials. **g**, **j**, Mean magazine entry probability (top) and mPFC Ca^2+^ activity (bottom) in response to cue for +RPE (**g**, yellow) or -RPE (**j**, cyan) trials and the trials (red) immediately following. **h**, Magazine entry during cue presentation was not affected by +RPE outcomes. **i**, Top: mPFC Ca^2+^ activity during cue presentation was not different between +RPE and subsequent trials. Bottom: Ca^2+^ decay was also not different between +RPE and subsequent trials. **k**, Magazine entries during cue presentation were significantly lower on trials that followed -RPE trials (paired t test, P= 0.0027). **l**, Similar to +RPE comparisons, mPFC Ca2+ rise (top) and decay (bottom) in response to cue presentation were not significantly different between -RPE trials the trials that followed. PSTHs are expressed as group mean ± shaded SEM. Before-after plots show both group mean ± SEM (black) and individual data as colored dots with lines (dotted lines=males; solid lines=females). \**P* < 0.05, \*\**P* < 0.01, \*\*\**P* < 0.001, \*\*\*\**P* < 0.0001; †*P* < 0.05, ††*P* < 0.01, †††*P* < 0.001, ††††*P* < 0.0001 from baseline.

As expected, the magnitude of reward seeking changed in response to unexpected appetitive outcomes with greater seeking during +RPE outcomes and less seeking during -RPE outcomes (**Figure 3d,e**). Consistent with the aversive task, appetitive outcomes induced a significant increase in mPFC Ca^2+^ response independent of value or RPE (**Figure 3d,f**). Interestingly, Ca^2+^ decay kinetics exhibited an inverse relationship to reward seeking such that a larger decay was observed following large reward delivery, when reward seeking was high, and a smaller decay was observed following reward omission, when reward seeking was lower than xRPE trials (**Figure 3d,f**). In contrast to the aversive task, neither reward seeking nor mPFC Ca^2+^ activity dynamically adjusted to unexpected appetitive outcomes in trial-by-trial comparisons (**Figure 3g-l**), although rewarding seeking was significantly decreased during cue presentation immediately following reward omission (**Figure 3j,k**).

Taken together, these data suggest that mPFC Ca^2+^ activity encodes the perceptual salience of both appetitive and aversive outcomes, as well as cues predictive of them, independent of valence, value, or outcome expectancy. Moreover, salience responsivity gradually shifts across conditioning from the distal to proximal cue (i.e., onset vs offset), in concert with the shift in motivational salience produced by associative learning. Interestingly, the observed cue-induced salience response adapted on a trial-by-trial basis in response to unexpected aversive, but not appetitive, outcomes. Lastly, the seemingly inverse relationship between behavioral responding and mPFC Ca^2+^ decay across learning and during RPE outcomes points to possible behavioral encoding by the mPFC.

### Distinct defensive phenotypes support movement encoding by the mPFC

To more closely investigate a possible role for the mPFC in encoding behavioral responding, we first took advantage of two distinct defensive strategies, freezing and darting, which emerged during aversive conditioning. Rats were classified *post hoc* as either darters or freezers based on their conditioned response to cue presentation (**Figure 4a-b** **and Supplemental Figure 2a-c**). In agreement with previous reports ^45^, females were more likely to exhibit conditioned darting than males (**Figure 4b**). Darters not only exhibited a greater degree of darting in response to aversive cue onset and offset, but also a higher velocity, more dart-like, response to shock delivery (**Figure 4c**). Accordingly, although both phenotypes exhibited more conditioned freezing in response to cue presentation across training, darters consistently exhibited less freezing than freezers throughout training and probe testing (**Supplemental Figure 4a-c and** **Figure 4d,e**). In turn, darters exhibited more movement during the cue presentation period and following shock delivery compared to freezers (**Supplemental Figure 4a,b,d and** **Figure 4d,f**). Interestingly, while both phenotypes exhibited comparable cue-induced increase in mPFC Ca^2+^, Ca^2+^ decay during cue presentation was significantly attenuated in darters compared to freezers, paralleling our earlier observations that freezing is inversely related to mPFC Ca^2+^ (**Figure 4d,g-h**). Likewise, while shock presentation increased mPFC Ca^2+^ activity in both freezers and darters, concomitant with shock-induced movement and reduced freezing, this effect was attenuated in darters, whose Ca^2+^ activity was already elevated likely constraining any additional increase due to a ceiling effect of GCaMP dynamics (**Supplemental Figure 4a,b,e and** **Figure 4d,g**).

To follow up on these findings, we next examined mPFC Ca^2+^ activity and movement patterns in relative isolation, without the confounds of perceptual and motivational salience, during both passive (**Figure 4i-k**, freezing onset) and active (**Figure 4l-n**, darting onset) defensive behavior. This analysis revealed a consistent decrease in mPFC Ca^2+^ at freezing onset when velocity also decreased (**Figure 4i**), and increased mPFC Ca2+ activity during darting onset when velocity increased (**Figure 4l**). Cross-correlational analyses further revealed a significant positive correlation between the leading velocity and lagging mPFC Ca^2+^ dynamics for both passive (**Figure 4j**) and active (**Figure 4m**) defensive behaviors. While darters exhibited slightly higher correlational strength between mPFC Ca^2+^ and velocity than freezers, the temporal dynamics did not differ significantly between phenotypes (**Figure 4k,n**). Taken together, these data suggest that the non-salience component of the mPFC Ca^2+^ dynamics during the aversive task is largely accounted for by differences in movement rather than encoding of a particular defensive strategy per se.

### mPFC Ca^2+^ activity and movement are positively correlated independent of behavioral valence

These findings led us to more closely examine the relationship between mPFC Ca^2+^ and movement across diverse and distinctly valenced behaviors. To distinguish the valenced, task-defined behaviors, such as defense and reward seeking, from generalized movement occurring regardless of task periods or behavioral types, we henceforth refer to the latter as task-uninstructed movement^50^, measured by continuous center-point velocity. Indeed, mPFC Ca^2+^ activity was positively correlated with velocity during both freezing onset (**Figure 5a-b**) and offset (**Figure 5d-e**). Interestingly, a similar relationship was observed in the appetitive task such that anticipatory reward seeking coincided with decreased velocity of movement and decreased mPFC Ca^2+^ activity (**Figure 5g-h**). Likewise, reward retrieval was associated with decreased velocity and positively correlated with mPFC Ca^2+^ activity (**Figure 5j-k**). Neither the correlational strength nor temporal delay differed across aversive (**Fig 5c,f**) or appetitive (**Fig 5i,l**) learning except for freezing onset (**Figure 5c**). These direct assessments of the relationship between mPFC Ca^2+^ activity and task-defined behavioral events suggests that the mPFC encodes uninstructed movement in a valence-independent manner. Consistently, re-examination of movement patterns during task events revealed that velocity captured changes in mPFC Ca^2+^ dynamics outside of salience encoding across all stages of the aversive and appetitive task (**Supplemental Figure 5**).

**Figure 4:**
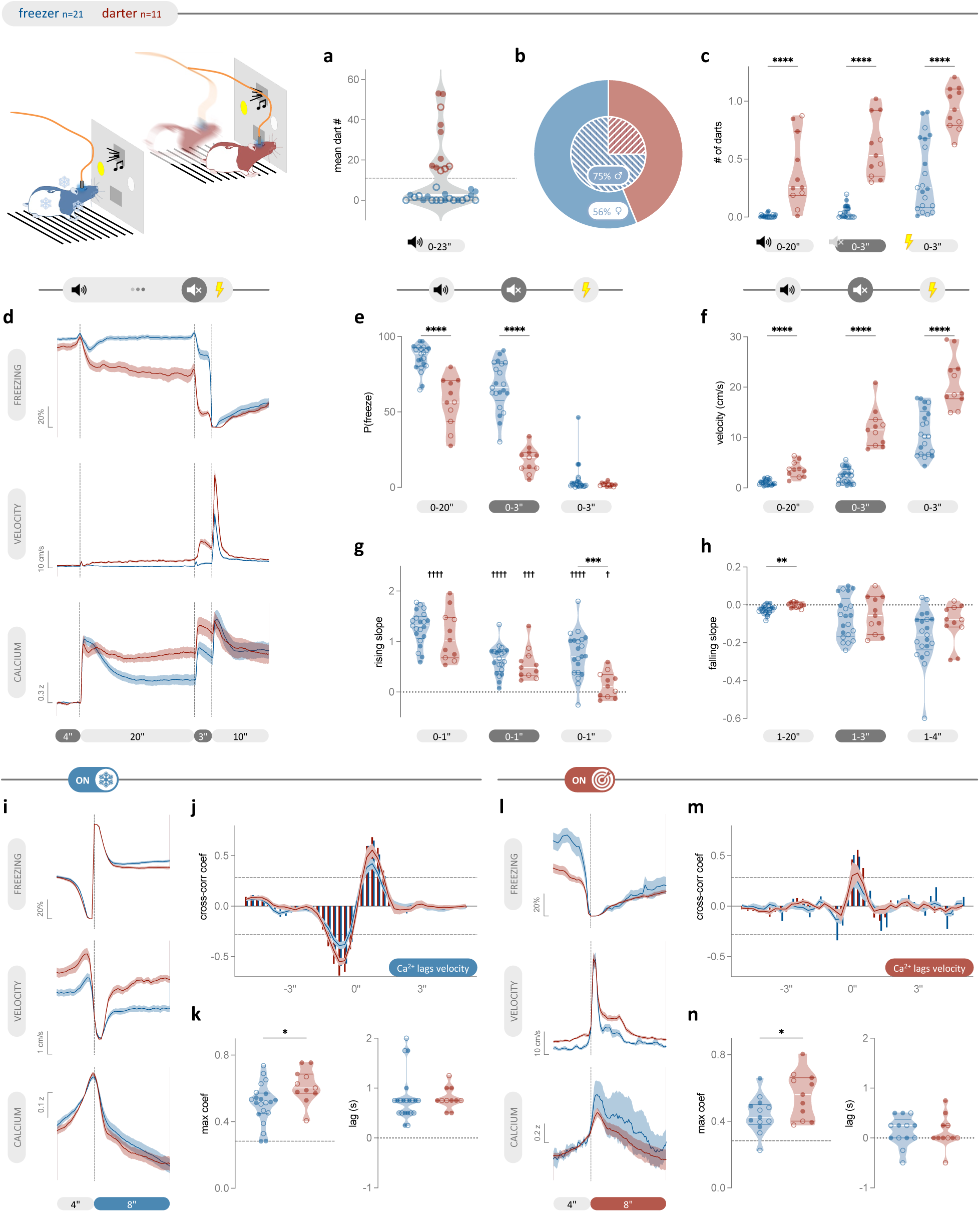
Distinct defensive phenotypes suggest that mPFC Ca^2+^ activity encodes movement. **a**, Rats classified as freezers exhibited little-to-no darts during cue presentation as exemplified by the violin plot showing dart counts per probe session. Dashed line indicates the threshold for phenotype classification. **b**, Darting was more prevalent in females than males. **c**, Darters exhibited more darting during cue periods and a high velocity, dart-like response to shock delivery (unpaired t test, P<0.0001). **d**, Mean freezing probability (top), velocity (middle), and mPFC Ca^2+^ activity (bottom) in response to aversive cue presentation and shock delivery during xRPE probe trials. **e-f**, Darters exhibited significantly less freezing during cue presentation (**e**, unpaired t test, P<0.0001), and higher velocity movement during both cue presentation and shock delivery (**f**, unpaired t test, P<0.0001). **g**, While both phenotypes showed increased mPFC Ca^2+^ activity from baseline in response to cue presentation (paired t test, cue onset: P<0.0001, cue offset: P<0.0001 for freezers and P=0.0002 for darters) and shock delivery (paired t test, P<0.0001 for freezers and P= 0.0366 for darters), Ca^2+^ response was greater in freezers than darters in response to shock (unpaired t test, P=0.0003). **h**, mPFC Ca^2+^ decay was significantly faster in freezers than darters during cue presentation (unpaired t test, 0.003). **i**, Mean freezing probability (top), velocity (middle), and mPFC Ca^2+^ activity (bottom) at freezing onset. **j**, Velocity and mPFC Ca^2+^ activity are significantly positively correlated with a (**k**) stronger relationship in darters than freezers (unpaired t test, P=0.0181) despite similar lag between the variables. **l**, Mean freezing probability (top), velocity (middle), and mPFC Ca^2+^ activity (bottom) at darting onset. **m**, Velocity and mPFC Ca2+ activity are significantly positively correlated. **n**, The strength of this correlation is higher in darters (unpaired t test, P= 0.0297) while the lag corresponding to the maximum positive cross-correlation does not differ between phenotypes. PSTHs are expressed as group mean ± shaded SEM. Violin plots indicate median and quartiles. Individual data are shown as circles (open=males; closed=females). Cross-correlograms show the cross-correlation function (XCF) between the group average velocity and mPFC Ca^2+^ activity (shown in bars), and the group average of individual rat’s XCF (shown as mean ± shaded SEM). Dashed lines on the cross-correlograms and corresponding max coef violin plots indicate significance cutoff. \**P* < 0.05, \*\**P* < 0.01, \*\*\**P* < 0.001, \*\*\*\**P* < 0.0001; †*P* < 0.05, †††*P* < 0.001, ††††*P* < 0.0001 from baseline.

**Figure 5:**
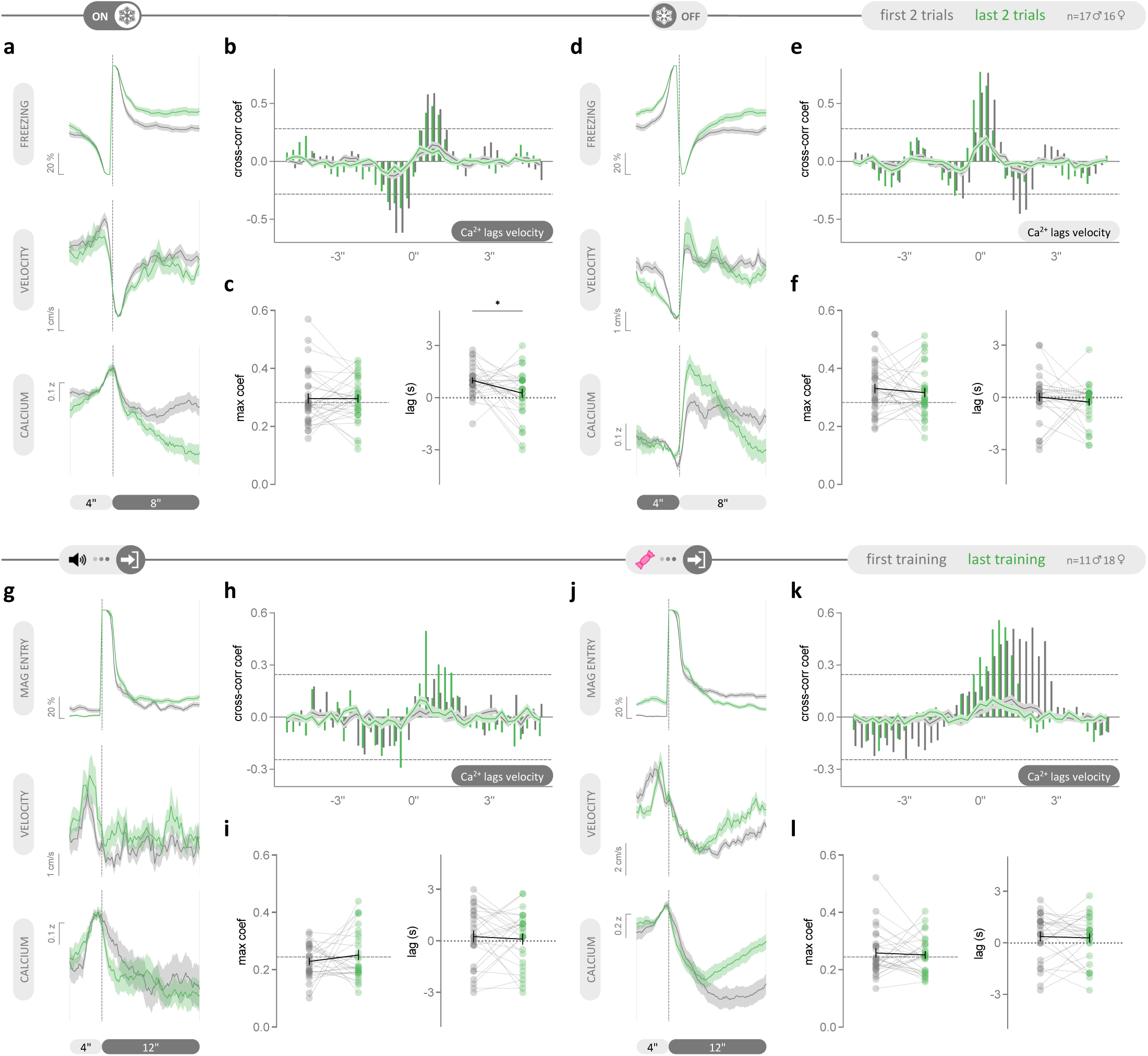
mPFC Ca^2+^ activity shows consistent, positive correlation with movement across distinct appetitive and defensive behaviors. **a**, **d**, Mean freezing probability (top), velocity (middle), and mPFC Ca^2+^ activity (bottom) at freezing onset (**a**) and offset (**d**) during the first (grey) and last two (green) conditioning trials. **b**, **e**, Velocity and Ca^2+^ activity are positively correlated with Ca^2+^ lagging both freezing onset (**b**) and offset (**e**). **c**, **f**, The strength of the positive cross-correlation between velocity and Ca^2+^ (left) and the corresponding lag (right) do not differ across training except for freezing onset (**c** right, paired t test, P= 0.0230). **g**, **j**, Mean magazine entry probability (top), velocity (middle), and mPFC Ca^2+^ activity (bottom) at the first magazine entry during cue presentation (**g**) and after pellet delivery (**j**) during the first (grey) and last (green) conditioning session. **h**, **k**, Velocity and Ca^2+^ activity are significantly positively correlated with Ca^2+^ lagging the first magazine entry during cue presentation (**h**) and after pellet delivery (**k**). **i**, **l**, The strength of the positive cross-correlation between velocity and Ca^2+^ (left) and the corresponding lag (right) do not differ across training. PSTHs are expressed as group mean ± shaded SEM. Before-after plots show both group mean ± SEM (black) and individual data as dots with lines (dotted lines=males, solid lines=females). Cross-correlograms show the cross-correlation function (XCF) between the group average velocity and Ca^2+^ activity (shown in bars) and the group average of individual rat’s XCF (shown as mean ± shaded SEM). Dashed lines on the cross-correlograms and before-after plots indicate significance cutoff.

### mPFC salience encoding supports adaptive reward seeking

In contrast to the distinct defensive phenotypes uncovered during aversive Pavlovian conditioning, there was no evidence of divergent response strategies in rats that underwent appetitive conditioning. However, individual differences in mPFC Ca^2+^ emerged over the course of testing in response to appetitive outcomes such that some rats exhibited a robust salience-like increase in Ca^2+^ activity in response to reward delivery, whereas others exhibited no such response. We took advantage of this natural randomization to investigate the potential functional significance of divergent salience encoding. To this end, rats were classified *post hoc* as pellet-responsive if they consistently exhibited increased mPFC Ca^2+^ activity in response to expected pellet delivery, or non-responsive if the decay in Ca^2+^ activity following cue offset was consistently unaffected by reward delivery (**Figure 6a-c**). The number of responsive and non-responsive rats was similar across sexes (**Figure 6b**). Comparison of mPFC Ca^2+^ activity between responsive and non-responsive rats found that while both phenotypes exhibited the aforementioned salience-induced increase in Ca^2+^ during cue presentation, the magnitude of this response was significantly more pronounced in the pellet responsive phenotype. Moreover, only the responsive phenotype exhibited a salience-like response to reward delivery (**Figure 6d-e**). Consequently, the subsequent decay in mPFC Ca^2+^ associated with reward retrieval and an overall decrease in movement (**Figure 6d,g-h**), was interrupted in the pellet-responsive phenotype (**Figure 6d,f**). These data suggest that a subset of rats exhibited heightened mPFC encoding of salience in response to both the conditioned stimulus and reward delivery.

**Figure 6:**
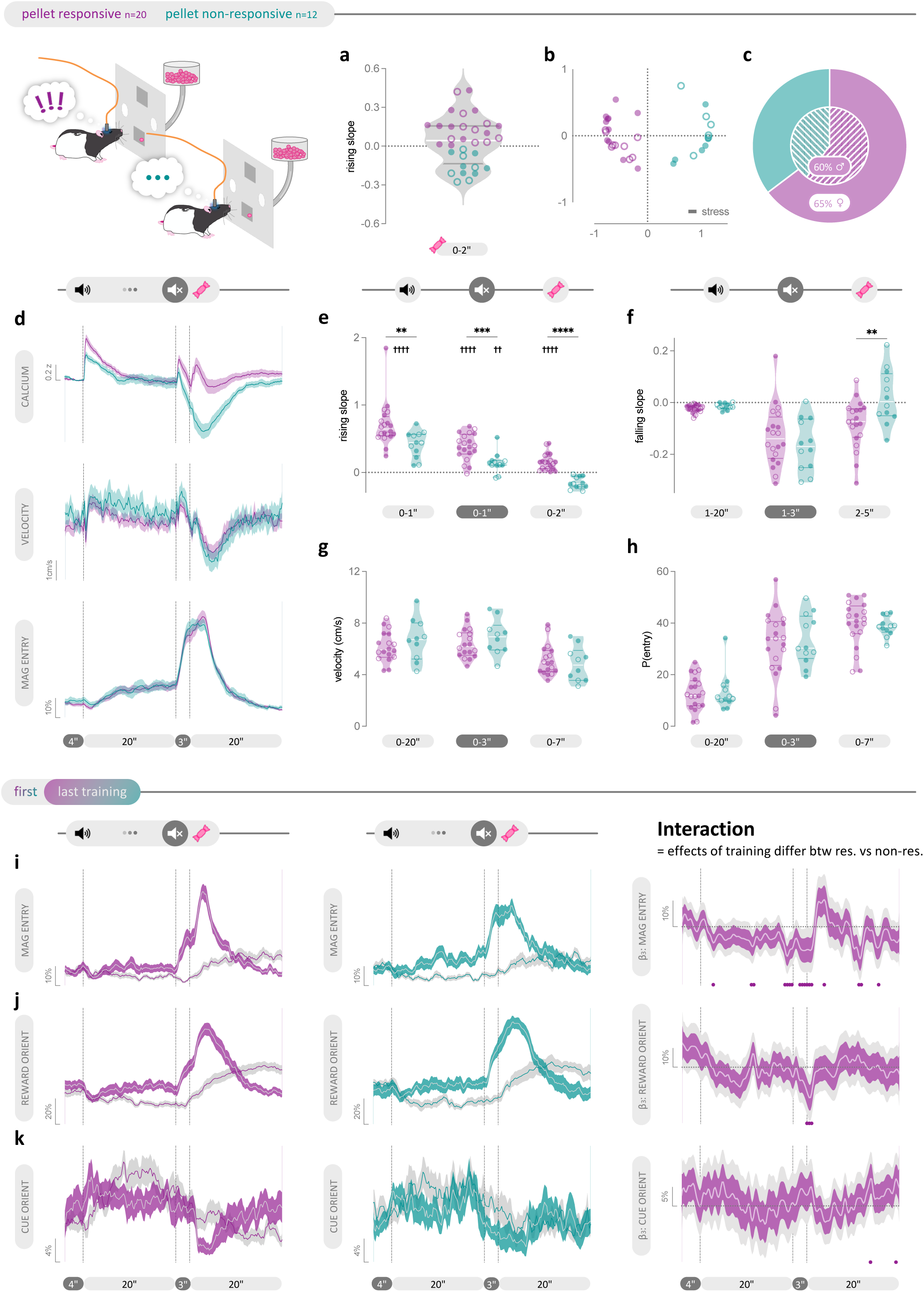
Distinct salience encoding supports adaptive reward seeking. **a**, Violin plot showing the distribution of the average rise in mPFC Ca^2+^ following pellet delivery during xRPE probe trials. Dotted line at y=0 indicates the threshold for phenotype classification. **b,** MDS visualization of RDM illustrating the dissimilarity in average pellet-induced Ca^2+^ activity across subjects and phenotypes. **c**, The prevalence of the pellet-responsive phenotype was similar between males and females. **d**, Mean mPFC Ca^2+^ activity (top), velocity (middle), and probability to enter the food magazine (bottom) in response to appetitive cue presentation and reward delivery during xRPE trials. **e**, While both phenotypes exhibited increased mPFC Ca^2+^ activity in response to cue presentation relative to baseline (paired t test, onset: P<0.0001, offset: P<0.0001 for responsive, P=0.0055 for non-responsive), the increase in the pellet-responsive phenotype was significantly larger than the non-responsive phenotype (unpaired t test, onset: P=0.0093, offset: P=0.001), and only the responsive phenotype exhibited a significant increase in Ca^2+^ from baseline in response to pellet delivery (paired t test, P<0.0001 for responsive; unpaired t test, P<0.0001). **f**, The Ca^2+^ decay following reward delivery was significantly shifted forward in time in the responsive vs nonresponsive phenotype (unpaired t test, P=0.0062). **g-h**, There were no significant differences in behavioral performance between phenotypes, including velocity (**g**) and magazine entry probability (**h**). **i**, Mean magazine entry probability in response to cue presentation and reward delivery during the first (grey shade) and last (color shade) conditioning session for the pellet-responsive (left) and non-responsive phenotype (middle). Functional coefficient estimates (right) showing that, across training, pellet-responsive rats have a smaller increase in reward seeking during cue presentation and the delay preceding reward delivery but a greater increase in reward seeking immediately after pellet delivery compared to the non-responsive phenotype. **j**, Mean reward-oriented behavior in response to cue presentation and reward delivery during the first (grey shade) and last (color shade) conditioning session for the pellet-responsive (left) and non-responsive phenotype (middle). Functional coefficient estimates (right) showing attenuated increase in reward-oriented behavior during pellet delivery in pellet responsive vs non-responsive rats across training. **k**, Mean cue-oriented behavior in response to cue presentation and reward delivery during the first (grey shade) and last (color shade) conditioning session for the pellet-responsive (left) and non-responsive phenotype (middle). Functional coefficient estimates (right) showing significantly greater increase in cue-oriented behavior during the ITI in pellet responsive vs non-responsive rats across learning. PSTH are expressed as group mean ± shaded SEM. Violin plots indicate median and quartiles. Individual data are shown as circles (open=males; closed for females). \*\**P* < 0.01, \*\*\**P* < 0.001, \*\*\*\**P* < 0.0001; ††*P* < 0.01, ††††*P* < 0.0001 from baseline. FLMM coefficients are expressed as estimates ± pointwise 95% CIs (dark shade) and joint 95% CIs (light shade) accounted for multiple comparisons. Statistical significance was accepted if the joint 95% CI of a timepoint did not contain y=0 (indicated by the horizontal dotted line) and was indicated by filled circles along the x axis.

Interestingly, reward seeking and overall velocity of movement were not different between phenotypes during either cue presentation or reward delivery (**Figure 6d,g-h**) suggesting that mPFC salience signaling is unaffected by, and thus, separable from movement encoding. While overall behavioral responding did not differ between appetitive phenotypes, we hypothesized that sensory stimuli associated with reward delivery acquired incentive salience in pellet-responsive rats to drive differences in cue-orienting behavior across phenotypes. To this end, we captured cue- and reward-oriented behavior, akin to sign-tracking and goal-tracking, respectively, using pose estimation and kinematics (**Supplemental Figure 2d-f**). This analysis found no differences in reward-oriented behavior (**Supplemental Figure 6a,c,e**), cue-oriented behavior (**Supplemental Figure 6b,d**), or a bias toward either (**Supplemental Figure 6f**) between pellet-responsive and non-responsive phenotypes during xRPE probe trials. Both phenotypes also showed comparable and consistent positive correlation between Ca^2+^ activity and uninstructed movement across various task-defined behavioral events, including anticipatory reward approach (**Supplemental Figure 7a-c**), reward retrieval (**Supplemental Figure 7d-f**), onset (**Supplemental Figure 7g-i**) and offset (**Supplemental Figure 7j-l**) of reward-oriented behavior, and onset (**Supplemental Figure 7m-o**) and offset (**Supplemental Figure 7p-r**) of cue-oriented behavior, further supporting non-specific movement encoding by the mPFC.

Suspecting a potential asymptotic effect on behavioral responding during probe trials, which occurred well after training, we re-examined behavior by phenotype during training. Seeking sufficient statistical power to undercover subtle phenotypic effects that could be masked by summary statistics, we leveraged functional linear mixed modeling (FLMM) to estimate the timepoint-specific effects of phenotype and training on Ca^2+^ and behavioral timeseries (**Supplemental Figure 8a**). Coefficient estimates of the interactive effects of phenotype and training reinforced the phenotypic difference in salience responsivity, such that across training, the pellet-responsive phenotype exhibited an increase in mPFC Ca^2+^ in response to pellet delivery compared to the non-responsive phenotype (**Supplemental Figure 8b**). Small variations in velocity were observed during cue presentation and ITI (**Supplemental Figure 8c**). Interestingly, reward seeking was significantly lower in pellet-responsive compared to non-responsive rats during cue presentation, pellet delivery, and ITI, but significantly greater immediately following pellet delivery (**Figure 6i**). These results suggest that the responsive phenotype strategically engaged in reward retrieval close to the delivery window instead of prematurely during cue presentation and ITI, possibly due to heightened incentive salience attributed to sensory stimuli associated with pellet delivery. Accordingly, the responsive phenotype also exhibited significantly less reward-oriented behavior at the time of pellet delivery (**Figure 6j**), and greater cue-oriented behavior during the ITI (**Figure 6k**) relative to the non-responsive phenotype.

In support of the idea that the responsive phenotype used environmental salience to guide adaptive reward seeking, FLMM showed that the pellet-responsive rats not only exhibited increased mPFC Ca^2+^ in response to pellet delivery during xRPE trials relative to omission during -RPE trials (**Supplemental Figure 8a-b**) without changes in movement (**Supplemental Figure 8c**), but also adaptively engaged in more reward seeking following pellet delivery (**Supplemental Figure 8d**) and less reward-oriented behavior during the ITI (**Supplemental Figure 8e**) compared to non-responsive rats. Trial-by-trial changes in cue-oriented behavior did not differ between phenotypes (**Supplemental Figure 8f**). Taken together, these data show that salience responsivity toward task relevant stimuli predicts adaptive rewarding seeking during appetitive conditioning and in response to reward omission.

### mPFC activity bidirectionally regulates movement

Collectively, our findings suggest that mPFC activity encodes a combinatorial sensorimotor signal during Pavlovian appetitive and aversive task performance. While mPFC activity has previously been associated with salience encoding^51–55^ as well as various motivated behaviors^31, 32, 56–69^, a direct, albeit indiscriminate role for mPFC in uninstructed motor performance is somewhat surprising. Thus, to dissect the causal relationship between region-wide mPFC activity, motivated behavior, and uninstructed movement, we next used *in vivo* optogenetics to bidirectionally manipulate pan-neuronal mPFC activity during behavioral responding while taking advantage of the dichotomous defensive phenotypes uncovered during aversive conditioning (**Figure 7a-c**). We hypothesized that channelrhodopsin (ChR2)-mediated optogenetic stimulation would bias the defensive strategy toward the active response, darting, whereas halorhodopsin (NpHR)-mediated optogenetic inhibition would bias the response toward passive freezing. Therefore, to maximize the chance of engendering defensive strategy reversal, we expressed ChR2 in male rats which are predominantly freezers and NpHR in female rats which have a higher incidence of darters (**Figure 4b**), with GFP controls comprised of both sexes. Following *ex vivo* validation of viral constructs (**Supplemental Figure 10a-b**), rats receiving viral injections and optic fiber implantation into the bilateral mPFC were trained on the aversive Pavlovian conditioning task (**Figure 7a-c**). FLMM was used to estimate the movement-dominant window of mPFC Ca^2+^ dynamics across trials (**Supplemental Figure 11a**) in order to develop light delivery parameters that selectively targeted movement-related activity while minimizing potential behavioral consequences of disrupting salience encoding. The model estimation of Ca^2+^ dynamics during average movement successfully recapitulated the empirical Ca^2+^ dynamics observed during aversive trials, confirming significant above-baseline salience response during cue and shock periods (**Supplemental Figure 11b**). Coefficient estimates revealed a significant correlation between mPFC Ca^2+^ and movement beginning 3 s after cue onset and lasting until shock delivery (**Supplemental Figure 11c-d**). Based on these findings combined with an understanding that GCaMP kinetics are significantly slower than action potential firing^70^, light delivery for optogenetic manipulation was initiated 2 s after cue onset and terminated immediately prior to shock delivery.

During baseline testing, 5 out of 8 ChR2 rats, 2 out of 7 NpHR rats, and 3 out of 6 GFP rats were identified as darters. During optogenetic probe testing, laser delivery had no effect on freezing or overall movement in GFP controls (**Figure 7d**) as expected. However, ChR2-mediated mPFC stimulation significantly reduced freezing during cue presentation (**Figure 7e,g**). Unexpectedly, a nonsignificant decrease in movement and darting behavior was also observed following cue offset in ChR2-expressing rats during stimulation trials (**Figure 7e,h**). Paradoxically, shock-induced startle (i.e., increased velocity) was significantly attenuated in ChR2-expressing rats during stimulation trials despite the fact that light delivery terminated prior to shock delivery (**Figure 7e,i**). In contrast, summary statistics found that behavior was unaffected by optogenetic mPFC inhibition (**Figure 7f,g-i**). Effects of optogenetic manipulation of either direction were consistent across defensive phenotypes.

Considering the subtle effect of optogenetic manipulation on behavior, we followed up with FLMM to more closely assess timepoint-level effects that could be masked by summary statistics (**Supplemental Figure 10c**). The model successfully recapitulated the behavioral performances of GFP-expressing controls at baseline, revealing significant freezing immediately prior to and during cue presentation as well as a significant increase in movement following cue offset and during shock presentation (**Supplemental Figure 10d**). Behavior was not significantly different between either ChR2- or NpHR-expressing rats and GFP-expressing controls on sessions without light delivery (**Supplemental Figure 10e-f**). In addition, coefficient estimates supported our original analysis showing that light delivery had no effect in GFP-expressing controls (**Figure 7j**). These estimates further confirmed that optogenetic mPFC stimulation reduces freezing and increases movement during cue presentation but decreases movement following cue offset and during shock delivery (**Figure 7k**). As was evident from the behavioral PSTHs, mPFC stimulation prevented adaptive behavioral responding to differentially salient cues whereby both freezing and movement appeared “clamped” at a steady state throughout the entire laser delivery period (**Figure 7e**). Conversely, optogenetic mPFC inhibition significantly reduced movement between cue offset and shock delivery as well as the ITI period (**Figure 7l**). Despite this, no significant effect of mPFC inhibition to increase freezing was observed during cue presentation (**Figure 7l**). However, this could easily be attributed to a ceiling effect. To address this, we also examined the effect of bidirectional optogenetic mPFC manipulation during the ITI period when threat responding was relatively low and during a separate session conducted in a non-threatening control context. Similar to our findings during cue presentation, mPFC stimulation during the ITI produced an immediate significant reduction in freezing and lasting increase in movement beyond the laser delivery period (**Figure 8a-b**). Consistently, light delivery in NpHR-expressing rats was associated with an immediate and lasting increase in freezing and decrease in movement relative to the period immediately preceding light delivery (**Figure 8a-b**). However, a similar, albeit more gradual pattern was also observed during light delivery in GFP-expressing controls suggesting that freezing naturally increases over time during the ITI as the onset of the next cue-shock presentation period approaches (**Figure 8a-b**). Importantly, however, rats in the GFP group maintained a consistent level of motor activity across time regardless of laser delivery when tested in a non-threatening control context, whereas optogenetic mPFC manipulation drove immediate and lasting bidirectional changes in movement in both ChR2-and NpHR-expressing rats (**Figure 8c-d**).

**Figure 7:**
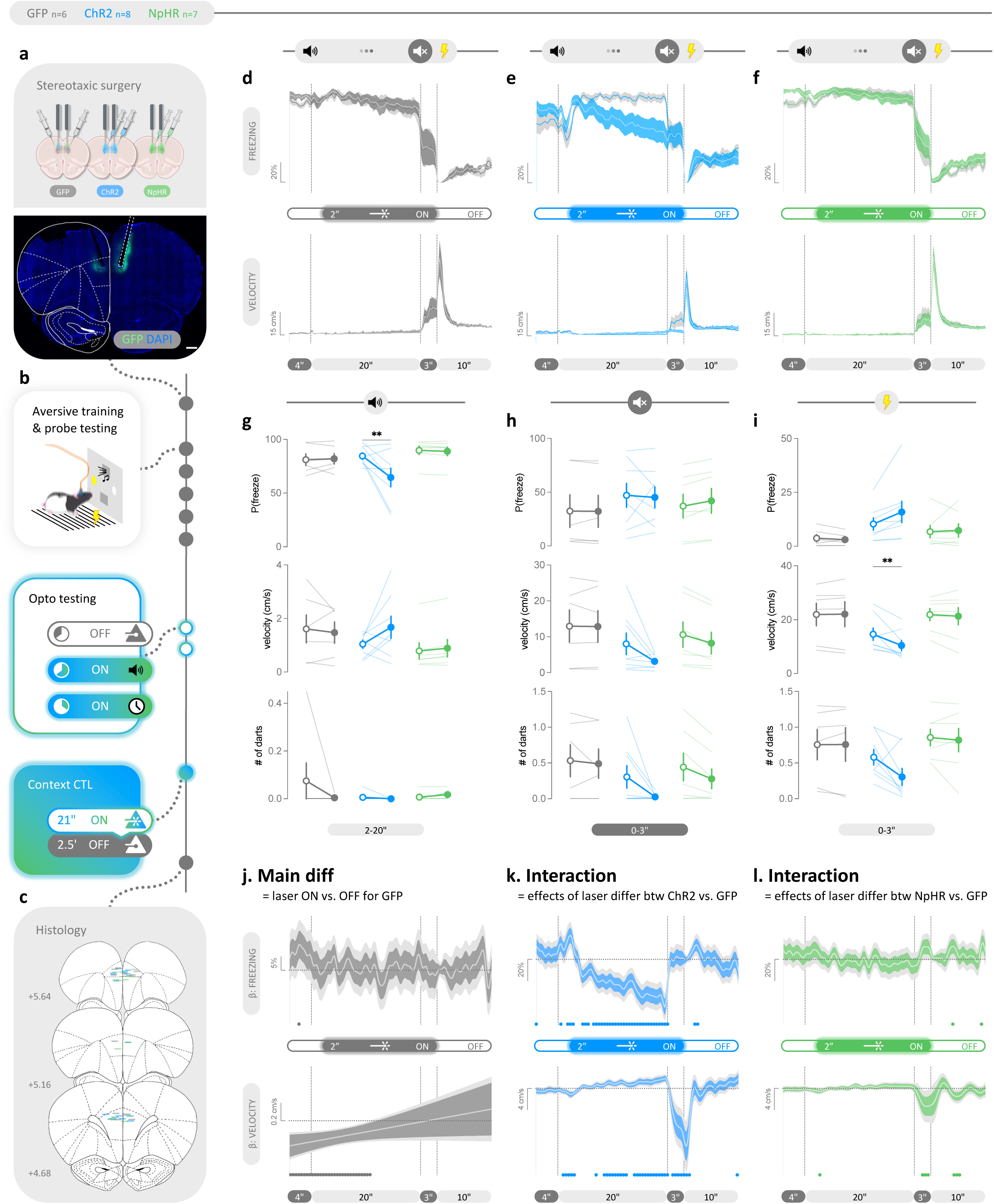
Bidirectional optogenetic manipulation of the mPFC during cue presentation disrupts movement regulation and adaptive threat responding. **a**, Top: Excitatory opsin (ChR2), inhibitory opsin (NpHR), or control fluorophore (GFP) was bilaterally expressed in the mPFC of male and female rats. Bottom: Representative expression and implant site localization. **b**, Following recovery, rats underwent the same aversive conditioning and testing procedure as outlined in Figure1b (top) followed by two optogenetic probe sessions (middle) and a context control session (bottom). **c**, Implant termination sites. **d-f**, Mean freezing probability (top) and velocity (bottom) in response to aversive cue presentation and shock delivery during laser OFF (light shade) vs. ON (dark shade) trials for rats expression GFP (**d**), ChR2 (**e**), and NpHR (**f**). **g-i**, Optogenetic stimulation of the mPFC during cue presentation significantly reduced average freezing probability during cue presentation (**g**, top, two-way RM ANOVA, interaction: P=0.0209; post hoc Šídák, ChR2: P=0.0028) and movement during shock delivery (**i**, middle, two-way RM ANOVA, interaction: P=0.0447; post hoc Šídák, ChR2: P=0.0052). Laser delivery reduced average movement (**h**, middle, two-way RM ANOVA, main effect of laser: P=0.0293) and dart counts (**h**, bottom, two-way RM ANOVA, main effect of laser: P=0.0226) during delay. **j-l**, Functional coefficient estimates showing (**j**) no meaningful effects of light delivery on behavior in GFP-expressing rats, (**k**) a significant reduction in freezing during cue presentation (top) concomitant with increased movement during cue, but reduced movement during the delay and shock period (bottom) in ChR2-expressing rats and (**l)** a significant reduction in movement during the delay period (bottom) in NpHR-expressing rats. PSTHs are expressed as group mean ± shaded SEM. Before-after plots show both group mean ± SEM (open=laser OFF; closed=laser ON) and individual data. FLMM coefficients are expressed as estimates ± pointwise 95% CIs (dark shade) and joint 95% CIs (light shade) accounted for multiple comparisons. Statistical significance was accepted if the joint 95% CI of a timepoint did not contain y=0 (indicated by the horizontal dotted line) and was indicated by filled circles along the x axis. \*\**P* < 0.01.

**Figure 8:**
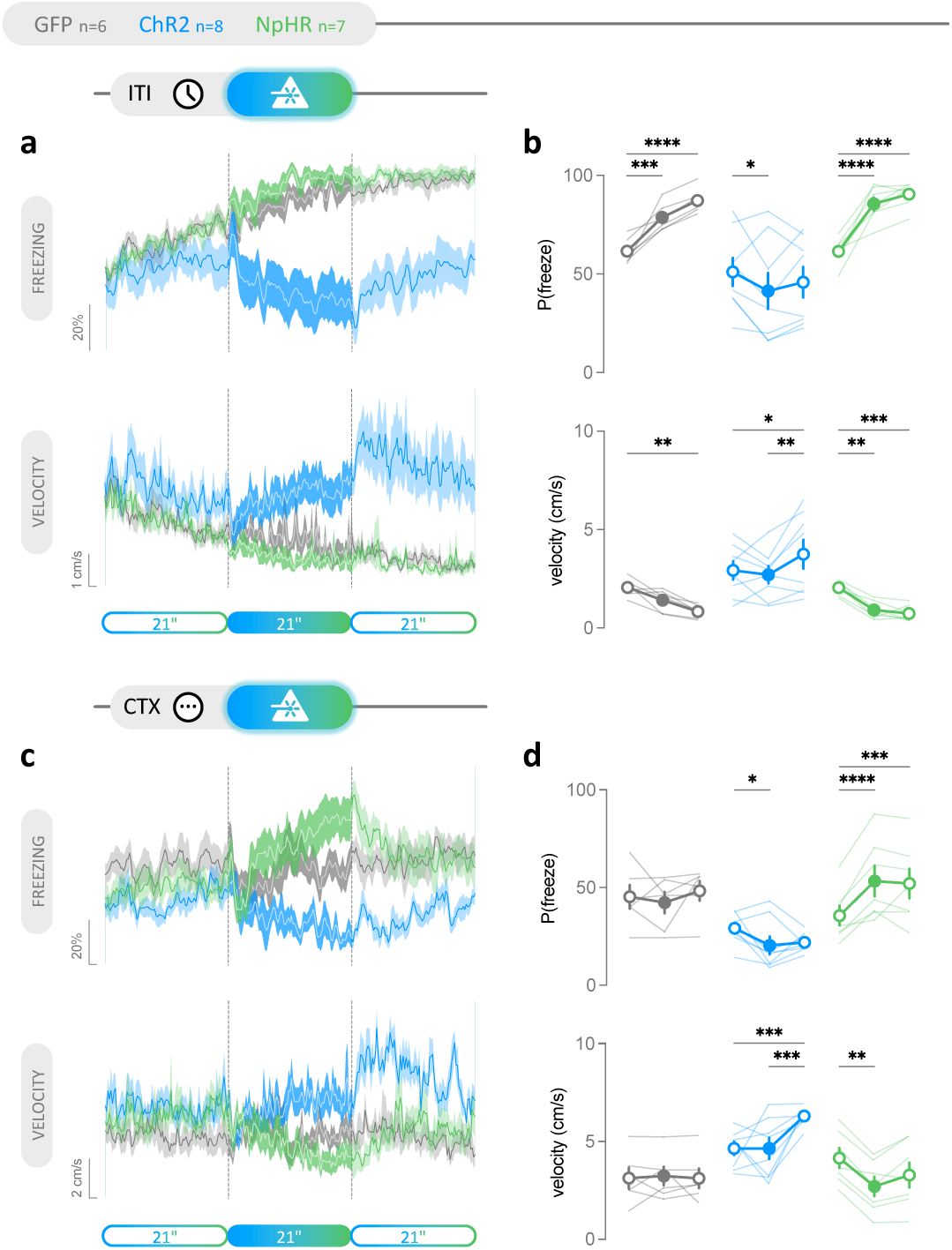
Optogenetic manipulation of mPFC activity bidirectionally drives immediate and lasting changes in movement. **a**, Mean freezing probability (top) and velocity (bottom) before (light shade), during (dark shade), and after (light shade) laser light delivery during the ITI of optogenetic probe trials. **b**, Optogenetic stimulation significantly reduced freezing (top, two-way RM ANOVA, interaction: P<0.0001; post hoc Tukey, ChR2: P=0.0312 for pre vs. ON) and produced a sustained increase in movement beyond the manipulation window (bottom, two-way RM ANOVA, interaction: P<0.0001; post hoc Tukey, ChR2: P=0.0017 for ON vs. post, P=0.0143 for pre vs. post). Light delivery was associated with increased freezing probability (top, post hoc Tukey, NpHR: P<0.0001 for pre vs. ON, P<0.0001 for pre vs. post) and reduced movement (bottom, post hoc Tukey, NpHR: P=0.0014 for pre vs. ON, P=0.0002 for pre vs. post) in both GFP- and NpHR-expressing rats. **c**, Mean freezing probability (top) and velocity (bottom) before (light shade), during (dark shade), and after (light shade) laser light delivery during a context control session. **d**, Optogenetic mPFC stimulation reduced freezing (top, two-way RM ANOVA, interaction: P<0.0001; post hoc Tukey, ChR2: P=0.0407 for pre vs. ON) and a produced sustained increase in movement beyond the manipulation window (bottom, two-way RM ANOVA, interaction: P=0.0001; post hoc Tukey, ChR2: P=0.0001 for ON vs. post, P=0.0001 for pre vs. post). Optogenetic mPFC inhibition resulted in an immediate and sustained increase in freezing (top, post hoc Tukey, NpHR: P<0.0001 for pre vs. ON, P=0.0002 for pre vs. post), and a reduction in movement (bottom, post hoc Tukey, NpHR: P=0.0014 for pre vs. ON). In contrast, laser light delivery had no effect on either freezing or movement in GFP-expressing controls. PSTHs are expressed as group mean ± shaded SEM. Before-after plots show both group mean ± SEM (open=laser OFF; closed=laser ON) and individual data. \**P* < 0.05, \*\**P* < 0.01, \*\*\**P* < 0.001, \*\*\*\**P* < 0.0001.

Together, results from optogenetic perturbation of mPFC activity during both threatening and non-threatening conditions suggest that while mPFC activity is sufficient to engender task-uninstructed movement, physiological and likely ensemble- and/or circuit-specific dynamics, which were disrupted by supraphysiological pan-neural activation, are still required for adaptive active threat responding such as darting. On the other hand, while mPFC activity did not prove to be necessary for movement initiation and defensive responding during threat, the graded suppression of movement that occurs as a function of environmental salience during mPFC inhibition suggests that mPFC activity plays a gain-like modulatory role, instead of a strict stop/go switch for movement or defined defensive behaviors, such that the top-down control it exerts is able to bias movement to a certain extent until being overcome by intense motivational drive for survival behaviors.

## Discussion

Using a combination of *in vivo* calcium imaging, multidimensional behavioral segmentation, regression-based computational modeling, and bidirectional optogenetic perturbation, we found that the bulk mPFC activity consists of a stimulus-induced short-latency salience signal that is reported indiscriminately of stimulus valence or value, followed by a slower dynamical component that not only correlates with, but also casually modulates valence- and task-agnostic movement.

### Weighted summation model of salience detection

Salience is frequently classified as either sensory-perceptual, novel-surprising, or incentive-motivational^2, 55^. Given that mPFC has long been implicated in native^51, 52, 54, 55^ as well as maladaptive salience processing^4, 53^, it is likely that the bulk salience signal observed in the current study reflects a latent process of weighted integration across various salience modalities, with sensory-perceptual being the dominant contributor, offset by novel-surprising and incentive-motivational. This is supported by evidence from the present study indicative of mPFC encoding of all three salience types. First, we observed an indiscriminate increase in bulk mPFC Ca^2+^ activity in response to cues and outcomes of varying novelty, valence, value, and PE with the magnitude of this activity appearing to track the sensory intensity of the stimulus (audiovisual cue onset > foot shock > cue offset > pellet sound). In contrast, outcome omission of either valence precluded sensory-perceptual salience detection and the associated increase in mPFC Ca^2+^ activity. Second, the uniform attenuation of cue-induced mPFC Ca^2+^ activity observed across both appetitive and aversive learning, likely reflects the expected gradual diminishment of cue novelty that occurs over repeated exposures. Finally, both the uniform potentiation of mPFC Ca^2+^ during cue offset across training and the bidirectional alteration in this activity observed following aversive PE trials likely reflect changes to the motivational salience associated with cue offset – the most proximal cue to the outcome – perhaps even at a trial-level timescale. Importantly, results preclude a simpler, pure sensory account of this activity. Previous work found that sensory-induced activity in the primary visual cortex undergoes multiplicative modulation by a motion-linked gain factor^71^. Whether this is also the case in the present study with respect to salience signaling in the mPFC is unclear as bulk Ca^2+^ sensing prevents us from definitively discriminating whether mPFC salience detection underwent multiplicative integration, perhaps via an upstream gain signal, or additive modulation, mediated by a dedicated subpopulation, or more than likely a combination of the two motifs.

Our data also revealed the presence of two distinct mPFC Ca^2+^ activity profiles in the appetitive task, with 60-65% of subjects exhibiting robust mPFC salience encoding in response to reward delivery. Notably, this pellet-responsive phenotype exhibited an augmented salience response to other task events, despite lacking obvious differences in task-defined behaviors or uninstructed movement compared to the non-responsive phenotype. Prior evidence has linked potentiated neural response to cues with increased cue-oriented, or sign-tracking, behavior^46, 72^ and further linked this behavior to deficits in impulse control often associated with substance use disorders and attention deficit hyperactivity disorder^73, 74^. With this in mind, we hypothesized that pellet-responsive rats would exhibit increased cue-oriented behavior relative to their non-responsive counterparts, in essence representing a neural signature of disease vulnerability. However, this was not the case. Instead, we found that pellet-responsive rats engaged in less premature and more strategic reward retrieval, likely by using the sound of the pellet dispenser or reward delivery as a sensory cue. Again, in the absence of any differences in task environment between pellet-responsive and -nonresponsive phenotypes, these results point to a phenotypical divergence in salience detection accompanied by attentional processes as opposed to differences in sensory perception. Whether this phenotype is protective or becomes maladaptive upon exposure to environmental insults (e.g., addictive drugs) is an important future consideration given that salience misattribution is linked to risk for the development of and/or barrier to the recovery from various neuropsychiatric diseases^47, 52, 53, 72–77^. Altogether, these findings link mPFC salience encoding with multi-dimensional appetitive behaviors that have strong potential to inform clinically relevant disease modeling exploring cue responsivity localized within and beyond the mPFC.

### Dynamic gain hypothesis of behavioral regulation

An extensive literature supports a role for the mPFC in both appetitive and aversive responding^31, 32, 56–69^. However, recent findings revealing brain-wide sensorimotor encoding^50, 71, 78–80^ have raised credible concerns regarding the strength of conclusions made by studies that omitted either continuous multi-dimensional movement monitoring or rigorous experimental and statistical movement control. In the present study, we continuously monitored both task-defined behaviors and uninstructed movement alongside bulk recordings and explicitly modeled the relationship between mPFC Ca^2+^ activity and behavioral timeseries during both pre-defined task periods and self-directed behavioral responding. We found that mPFC bulk activity closely tracked generalized movement outside of salience encoding in a consistent, positive manner. This was the case across paradigms and was independent of learning, trial outcome, or motivated behaviors.

With these findings in mind, we hypothesized that mPFC activity bidirectionally regulates movement. Based on the differences in mPFC Ca^2+^ activity we observed between defensive phenotypes, we further hypothesized that increasing mPFC activity would facilitate a transition from passive (i.e., freezing) to active (i.e., darting) threat responding. Instead, we found that optogenetic perturbation of mPFC activity in either direction decreased movement velocity (i.e., suppressed darting) during cue offset – the trial period when darting is most frequently observed. Thus, while our original hypothesis was largely validated by our optogenetics findings, in combination with our photometry results, they suggest that the mPFC participates in the dynamic movement gain control network. Accordingly, optogenetic inhibition of the mPFC, a known association cortex, likely disrupted both the feed-forward motor efference during initial movement issurance and feed-back sensory reafference for ongoing movement modulation to varying extent. Its pecise gain function remains unresolved due to the kinetical limitation of the bulk recording modality employed here, which inevitably gives rise to Ca^2+^ activity that lags relative to movement onset, as well as the supraphysiological nature of pan-neuronal synchronous manipulation. This results in global suppression of movement as we observed in our context control experiment. Importantly, however, loss of gain does not necessarily translate to increased gating of afferent activity. Therefore, when movement instruction is sufficiently strong, presumably via amplification by cue- and context-dependent motivational processes as is the case in the aversive task, the absence of mPFC gain resulting from optogenetic inhibition can be overcome to facilitate both cue- and shock-elicited active threat responses (i.e., darting). On the other hand, we found that mPFC stimulation drove a non-specific increase in uninstructed movement rather than darting per se. This suggests that disruption of the native dynamic gain not only prevented the faithful amplification and transmission of defined sensorimotor signals but also scrambled or buried it under increased noise. This finding mirrors previous work showing improved mPFC signal-to-noise during aversive processing as a result of dopamine release^65^. In doing so, the diminished signal-to-noise prevented both active and passive adaptive threat responding.

Previous work has found that mPFC population activity contains latent dynamics orthogonal to those closely tracking sensory and movement during active avoidance, thus more credibly supporting its engagement in abstract cognition during fear states^69^. If cognitive / internal processes underlying defensive strategy or fear state indeed exist in latent population subspace, the dynamic gain hypothesis of mPFC behavioral regulation is the most parsimonious and defensible interpretation of our data. Rather than demonstrating a simultaneously necessary and sufficient role of the mPFC in mediating active vs passive fear response as has been suggested previously^57, 64, 67, 81^ or an innate defensive endophenotype^45^, our findings suggest that the dynamic, physiological activity of the mPFC is necessary for valence-agnostic behavioral responding. While we propose dynamic gain as a broad functional description to model the relationship between bulk mPFC neural activity and behavior, it remains agnostic of the underlying computational processes at the circuit-, ensemble-, and population-level that may carry out the specific gain function^58, 59, 61–64, 66–69, 82^.

### Functional bipartition model

Taken together, we propose a salience-movement bipartition schema for the region-wide mPFC dynamics, where a low-latency prominence broadcasts the integrated environmental salience perhaps as a global alert signal, while its ongoing activity acts as an online dynamic gain within the broader movement network to execute, monitor and regulate behavioral responding. A similar functional bipartition model has been proposed for the midbrain dopamine system to reconcile the low-latency, seemingly indiscriminate, valence- and value-free, salience-like phasic activity, and its canonical reinforcement learning function attributed to the slower component of dopaminergic activity, which seems to better correlate with task parameters such as value and prediction error^2, 83^. However, more recent work found that stimulus-induced salience signaling in dopamine neurons reliably predicted movement initiation, whereas its second component not only tightly correlates with movement but appears neither necessary nor sufficient for learning^84^. These findings cast doubt not only on the explanatory parsimony of putative dopamine-mediated error-based learning, but also perhaps its ontological validity.

In a similar vein, prior understandings regarding the mPFC’s role in error-related cognition largely rest on region-wide bulk activity with little or limited behavioral confound monitoring^11, 16, 18, 19, 24, 25, 27–30^, whereas non-specific bulk calcium imaging has been increasingly used to demonstrate brain-wide cognitive functions ^86–91^. In contrast, we did not observe a link between bulk mPFC Ca^2+^ dynamics and prediction error signaling not sufficiently accounted for by the salience-movement model. Indeed, fluctuations in Ca^2+^ signal were fully explained by integrated salience and general uninstructed movement across all stages of learning regardless of task-defined behavioral performance. Similar brain-wide sensorimotor multiplexing has been reported both at the region and single-cell level^2, 50, 67, 69, 71, 78, 79, 85, 92–95^ and was shown to follow an additive process in primary visual cortex^71^. Likewise, our results suggest that mPFC bulk activity is a combinatorial signal reflective of an additive integration between the salience and movement components. This was evident in the present study not only with sophisticated trial-level analyses using FLMM, but also remained appreciable after data were averaged across trials using conventional summary statistics.

### Sensorimotor is fundamental, cognition is emergent

Findings from the present study join recent work^67, 85, 96, 97^ to provide additional disconfirming evidence against the idea that cognitive / internal processes are directly observable from one-dimensional bulk or fixed weighted average neural activity where sensorimotor-related signals overwhelmingly predominate^50, 67, 71, 78–80, 85, 96^, or can be indirectly inferred from non-representative sampling of single-cell activity where mixed selectivity and multiplexing are the norm^22, 50, 71, 82, 92–95, 98^. Even if task-relevant variables can be readily decoded from high-dimensional population representation or low-dimensional latent dynamics^54, 67, 82, 92, 95, 98, 99^, without explicit experimental and/or statistical control for potential sensory-movement confounds, studies making positive inferences regarding cognitive processes have the potential to produce misleading conclusions. Recent evidence demonstrating population subspace encoding of movement-free, cognitive / internal processes^69, 96^ lends further credence to the idea that cognitive processes are “emergent” phenomena discernible only at the population-level ^93, 100, 101^.

That sensorimotor signaling appears distributed across regions ascribed complex cognitive functions like the mPFC likely has its origins in evolution. While mammals’ large brain size is thought to afford the increased cognitive capacity and behavioral flexibility necessary for conquering myriad diverse and complex ecological niches^102–104^, fossil records indicate that mammalian encephalization not only lagged body expansion^102^, but also began first with subcortical expansion tied to body control and locomotion, prior to the neocortical expansion that allowed for sensorimotor enhancement and integration^102, 103, 105^. As a result, the expanded frontier of possible neural state space likely laid most, if not all, prerequisite groundwork for the emergence of new cognitive processes and behavioral states^100, 106^. In other words, cognitive functions of increasing complexity and abstraction as emergent properties likely arose primarily through exaptation and/or as a byproduct of scaling of pre-existing cell types and wiring motifs, rather than necessitating entirely new classes of cells or structures.

In conclusion, our findings support a role for mPFC as an associative hub integrating externally driven salience detection and internally driven behavioral modulation across motivationally salient, yet distinctly valenced contexts. These results inform future translational work seeking to understand how mPFC impairment contributes to pathological states characterized by salience misattribution and behavioral dysregulation. Lastly, our work again stresses the importance of unbiased experimental and statistical movement control for *in vivo* experiments, in order to avoid spurious, however enticing, conclusions.

## DATA AVAILABILITY

Data have not been deposited in a public repository but will be available upon request. Custom code used is available on lab Github (<u>github.com/ejgloverlab</u>).

## ACKNOWLEDGMENTS

The authors thank Nathaly Arce Soto, Brianna Irizarry, Dianni N Kinarasri, Shannon Kolofske, Joseph Pitock, Kathryn Przybysz, Jacqueline Sanchez, Shree Srinivasan, and Hyerim Yang and for technical assistance. The authors appreciate thoughtful feedback during the Women in Alcohol Research Writing Retreat. This work was supported by NIH grants R01 AA031003, R01 AA029130, and R00 AA024208 to EJG.

## AUTHOR CONTRIBUTIONS

SH – experimental design, data collection, data analysis, manuscript preparation; LAR – data collection, data analysis, manuscript editing; EHM – data collection, data analysis, manuscript editing; JW – data analysis, manuscript editing; MNG – data analysis, manuscript editing; HW – data analysis, manuscript editing; EJG – experimental design, data analysis, manuscript preparation.

---

**Supplemental Figure 1:**
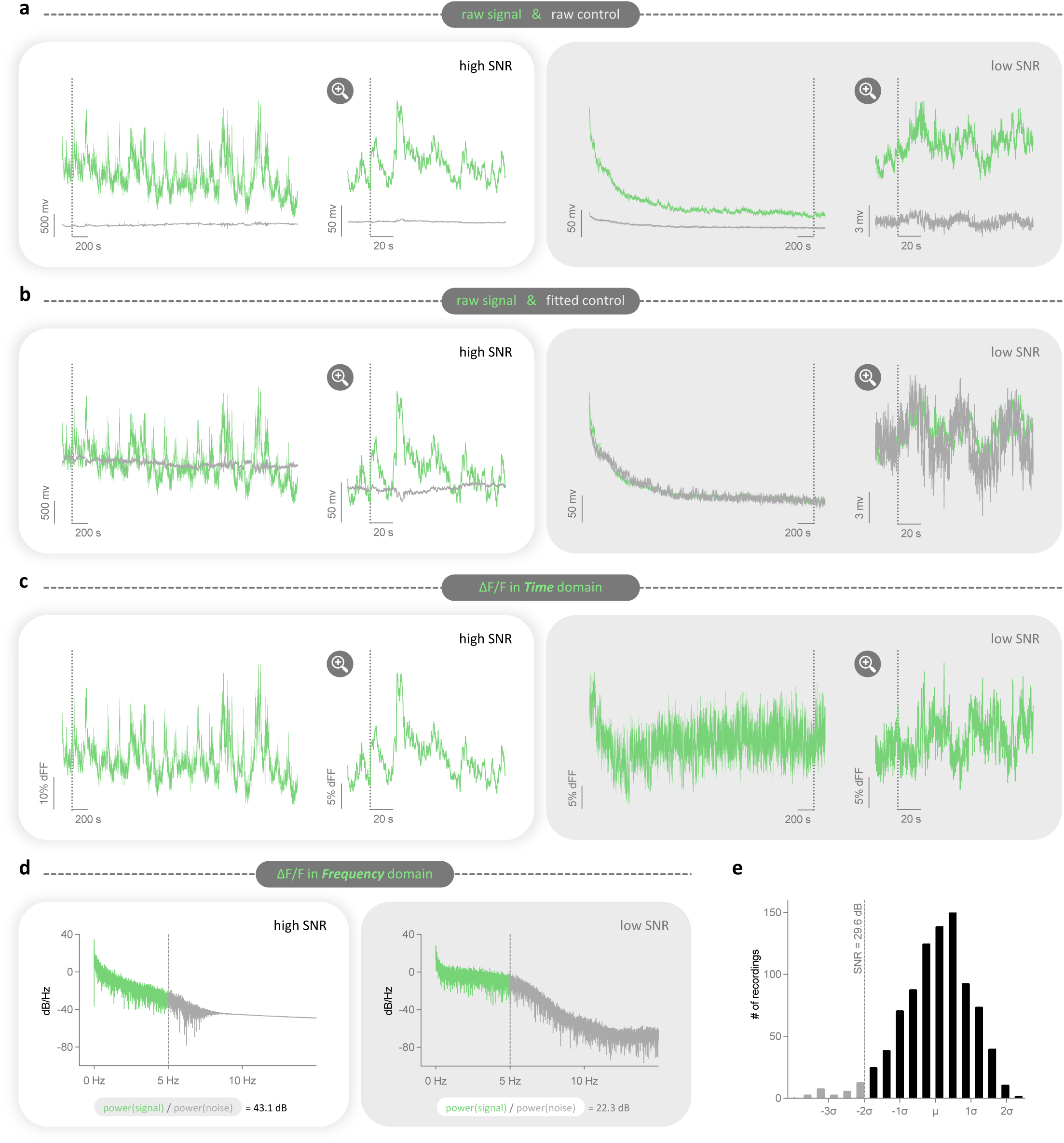
Photometry data pre-processing, noise-level quantification, and exclusion criterion. **a**, A representative raw Ca^2+^-dependent GCaMP signal trace (green) and isosbestic control trace (grey) of a high SNR recording (white panel) and a low SNR recording (grey panel). The session-wide trace (left) and a close-up of the same trace (right) aligned at the dotted line. **b**, The same trace as in (a) but with the isosbestic control (grey) fitted to the signal (green). **c**, Ca^2+^ timeseries expressed in percent ΔF/F. **d**, Periodograms showing the power spectral density estimate of ΔF/F, and the calculation of SNR from the average power of signal (green) vs. noise (grey) band. **e**, Distribution of SNRs from 891 recordings. Dashed line indicates the cutoff for exclusion, which was selected at two standard deviations below the mean.

**Supplemental Figure 2:**
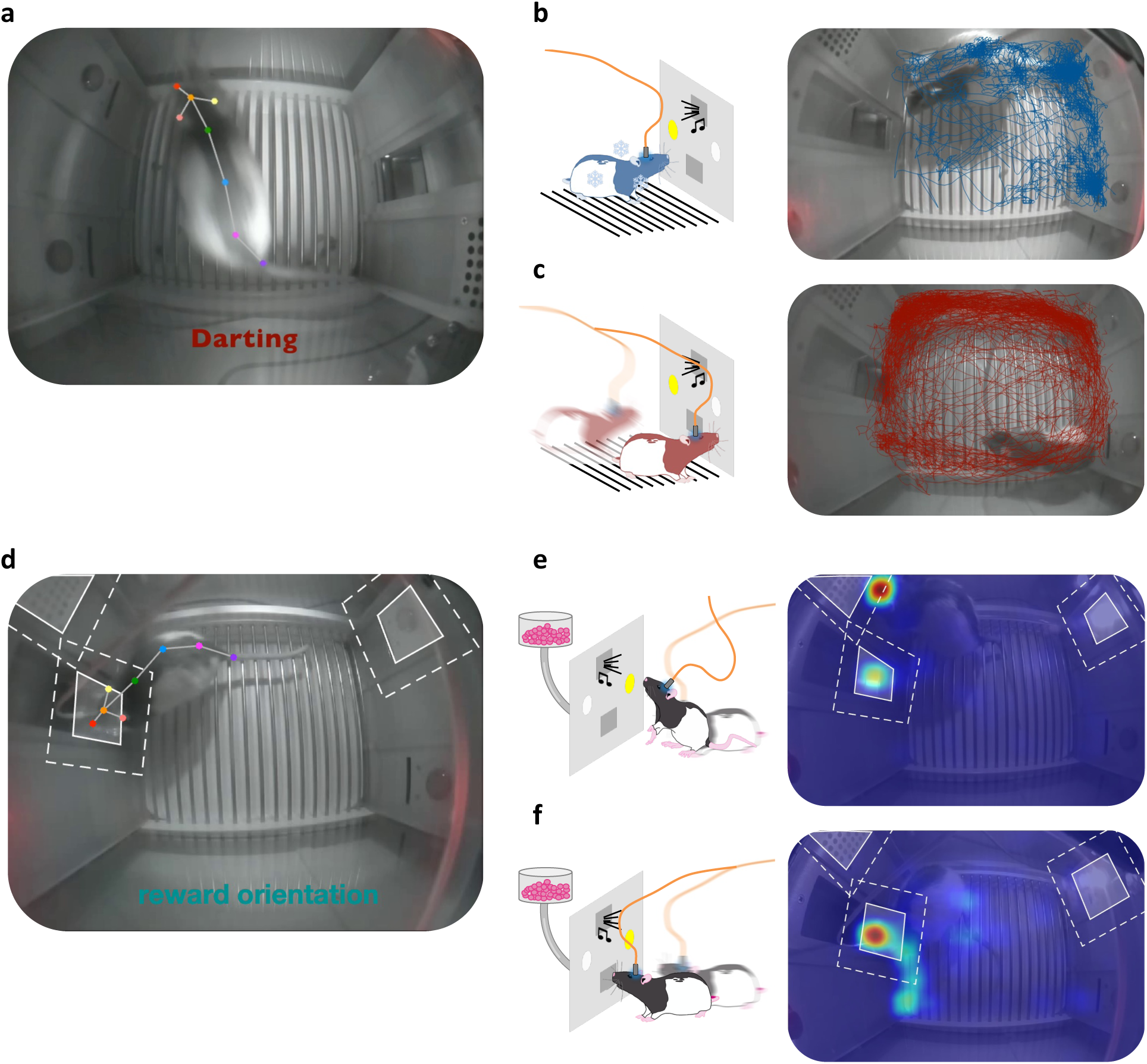
Pose estimation and kinematics for behavioral phenotyping. **a**, Frame capture of representative darting behavior (from Supplementary Movie 3) with keypoint and skeleton overlay. **b-c**, Representative center tracking trajectory of a freezer (**b**) vs. a darter (**c**) during an aversive probe session. **d**, Representative reward-oriented behavior (from Supplementary Movie 4) with keypoint, skeleton, and ROI overlay. **e-f**, Representative head location heatmap of a rat exhibiting significant (**e**) vs. relatively little (**f**) cue-oriented behavior during an appetitive probe session.

**Supplemental Figure 3:**
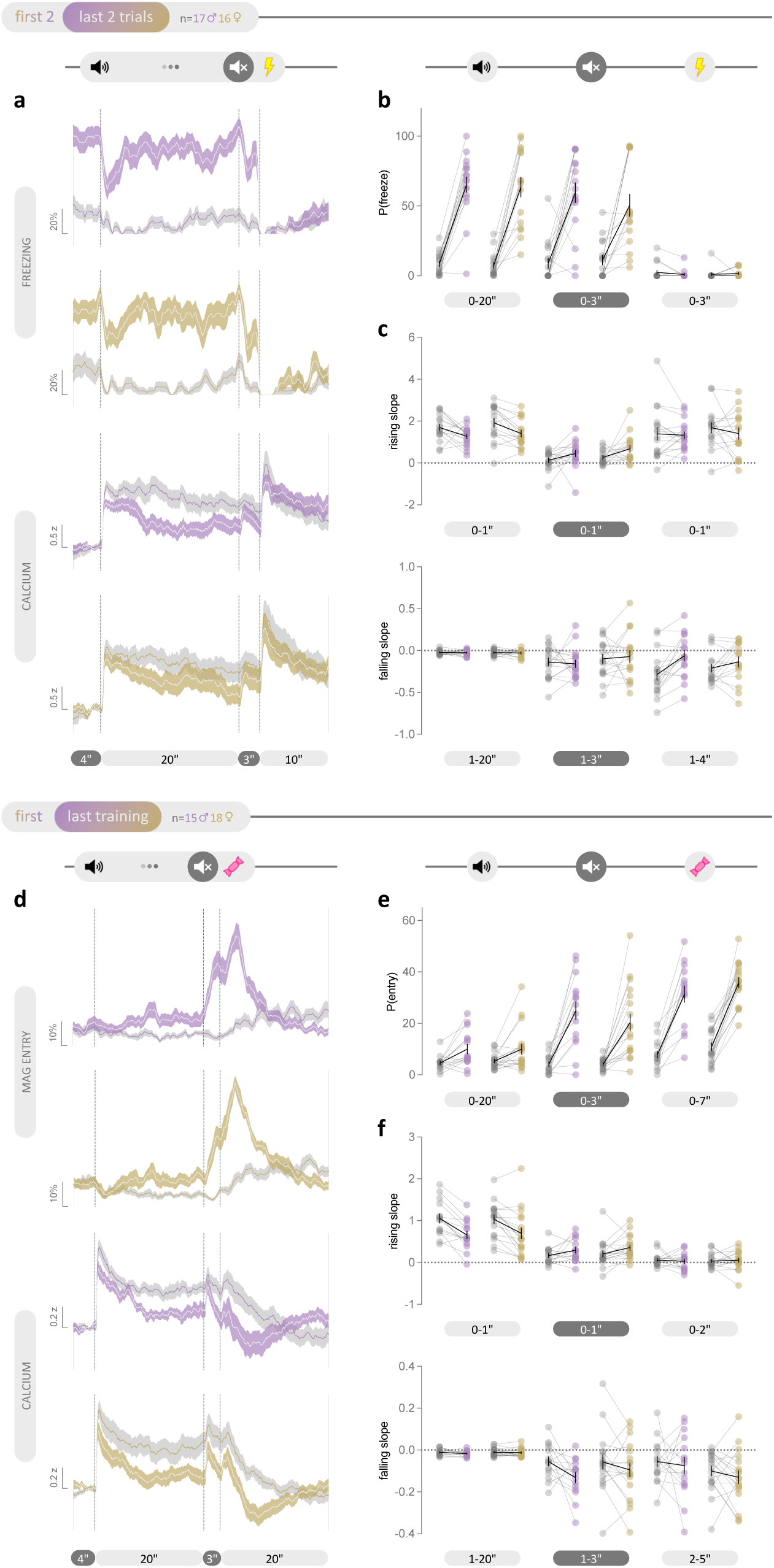
Absence of sex differences in mPFC Ca^2+^ activity or behavioral performance during either appetitive or aversive training. a, Mean freezing probability (top) and mPFC Ca^2+^ activity (bottom) in response to aversive cue and shock during the first (grey shade) and last two trials (color shade) of the aversive conditioning session. b-c, No sex difference was observed for freezing probability (b), or mPFC Ca^2+^ activity (c). d, Mean magazine entry probability (top) and mPFC Ca^2+^ activity (bottom) in response to cue and pellet delivery during the first (grey shade) and last appetitive conditioning session (color shade). e-f, No sex difference was observed for magazine entry probability (e), or Ca^2+^ activity (f). PSTHs are expressed as group mean ± shaded SEM. Before-after plots show both group mean ± SEM and individual data as dots with lines (purple=males; gold=females).

**Supplemental Figure 4:**
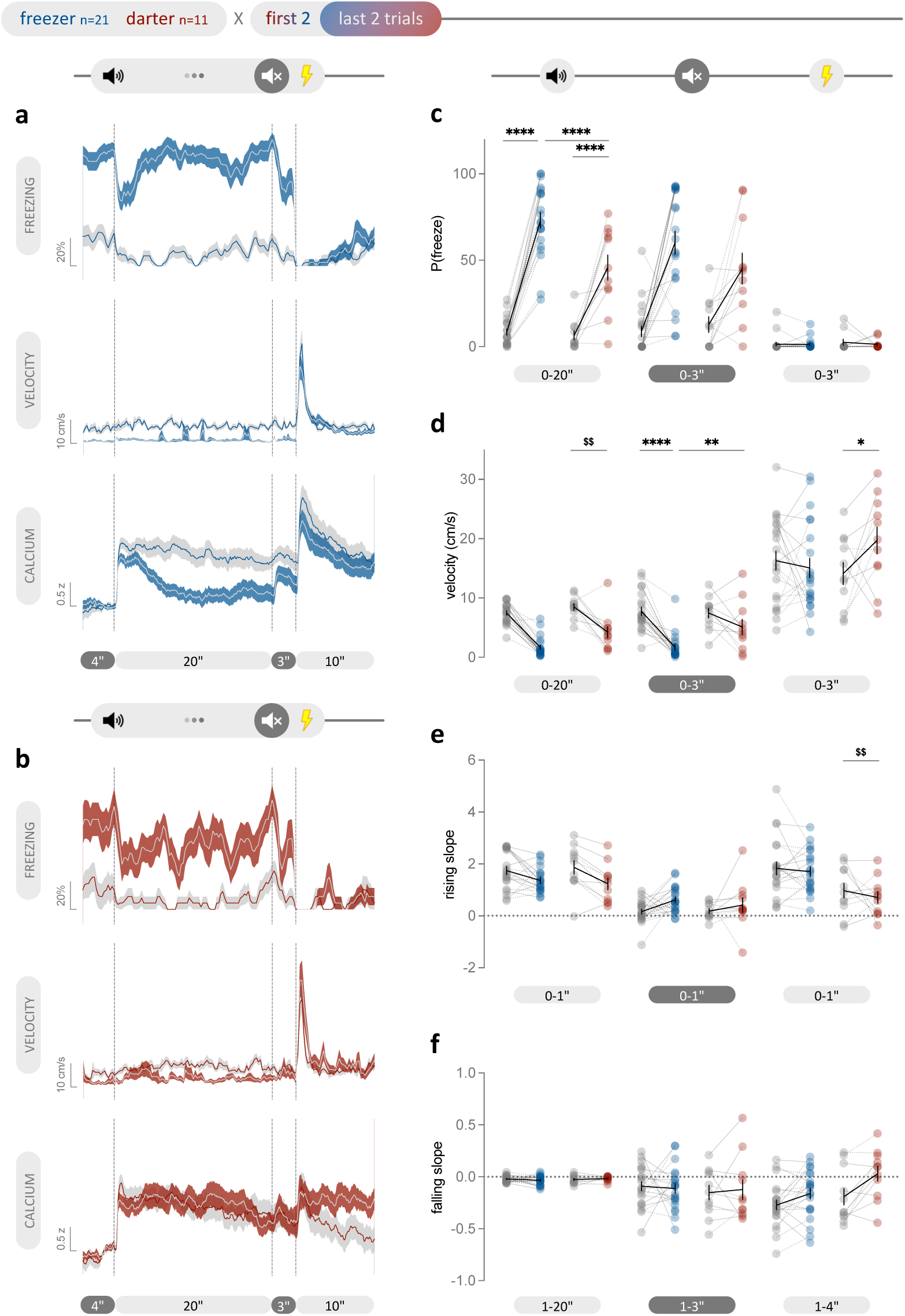
**Freezers & darters differ in mPFC Ca^2+^ activity and behavioral performance during training. a-b**, Mean freezing probability (top), velocity (middle), and mPFC Ca^2+^ activity (bottom) in response to aversive cue and shock during the first (grey shade) and last two trials (color shade) of the aversive conditioning session in freezers (**a**) and darters (**b**). **c**, Freezers exhibited a larger increase in freezing during cue presentation across training compared to darters (two-way RM ANOVA, interaction: P=0.0052, post hoc Fisher’s LSD, last: P<0.0001 for freezer vs. darter). **d**, Freezers exhibited less movement during cue presentation across training and as compared to darters (cue, two-way RM ANOVA, phenotype: P=0.0043; delay, two-way RM ANOVA, interaction: P=0.0192, post hoc Fisher’s LSD, freezer: P<0.0001 for first vs. last, last: P=0.0041 for freezer vs. darter). Shock response was potentiated in darters as evidenced by increased velocity across training (two-way RM ANOVA, interaction: P=0.0435, post hoc Fisher’s LSD, darter: P=0.0409). **e-f**, Shock-induced increase in mPFC Ca^2+^ activity was attenuated in darters than freezers throughout testing (**e**, two-way RM ANOVA, phenotype: P=0.0037). PSTHs are expressed as group mean ± shaded SEM. Before-after plots show both group mean ± SEM and individual data as dots with lines (dotted lines = males, straight lines = females). \**P* < 0.05, \*\**P* < 0.01, \*\*\*\**P* < 0.0001; ^$$^*P* < 0.01 main effect of phenotype.

**Supplemental Figure 5.**
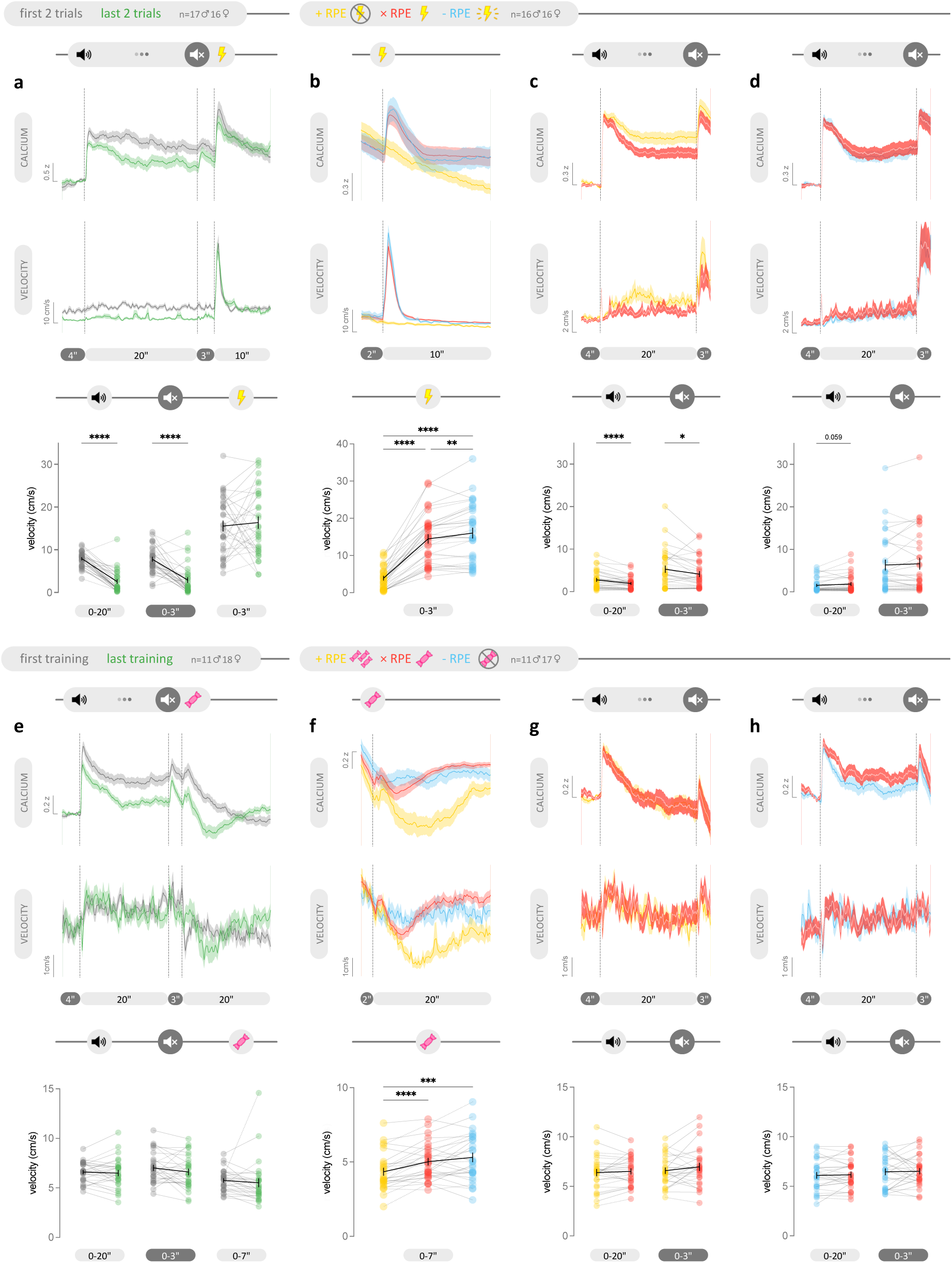
Changes in mPFC Ca^2+^ activity during appetitive and aversive tasks can be partially accounted for by changes in movement. a, Mean mPFC Ca^2+^ activity (top) and velocity (middle) in response to cue and shock during the first (grey) and last two trials (color) of aversive conditioning session showing decreased movement over learning during the shock-predicting cue (bottom, paired t test, P<0.0001). **b**, Mean mPFC Ca^2+^ activity (top) and velocity (middle) in response to expected mild shock (xRPE in red), unexpected shock omission (+RPE in yellow), and unexpected strong shock (-RPE in cyan), showing decreased movement during shock omission (bottom, one-way RM ANOVA, P<0.0001; post hoc Tukey, P<0.0001) and increased shock response to strong shock (post hoc Tukey, P=0.0032). **c-d**, Mean mPFC Ca^2+^ activity (top) and velocity (middle) in response to cue presentation during +RPE (**c**, yellow) or -RPE (**d**, cyan) trials, and the xRPE trials (red) that followed, showing decreased movement after shock omission (**c**, bottom, paired t test, cue: P<0.0001, delay: P=0.0101) but increased movement after strong shock (**d**, bottom, paired t test, cue: P=0.0587). **e**, Mean mPFC Ca^2+^ activity (top) and velocity (middle) in response to cue presentation and pellet delivery during the first (grey) and last conditioning session (green). **f**, Mean mPFC Ca^2+^ activity (top) and velocity (middle) in response to expected reward (xRPE in red), unexpected larger reward (+RPE in yellow), and unexpected reward omission (-RPE in cyan), showing decreased movement after large reward (bottom, mixed-effects analysis, P=0.0001; post hoc Tukey, P<0.0001). **g-h**, Mean mPFC Ca^2+^ activity (top) and velocity (middle) in response to cue presentation during +RPE (**g**, yellow) or -RPE (**h**, cyan) trials, and the xRPE trials (red) that followed showing no trial-by-trial changes in velocity (bottom). PSTHs are expressed as group mean ± shaded SEM. Before-after plots show both group mean ± SEM and individual data as dots with lines (dotted lines = males, straight lines = females). \**_P_* < 0.05, \*\**_P_* < 0.01, \*\*\**_P_* < 0.001, \*\*\*\**_P_* < 0.0001.

**Supplemental Figure 6:**
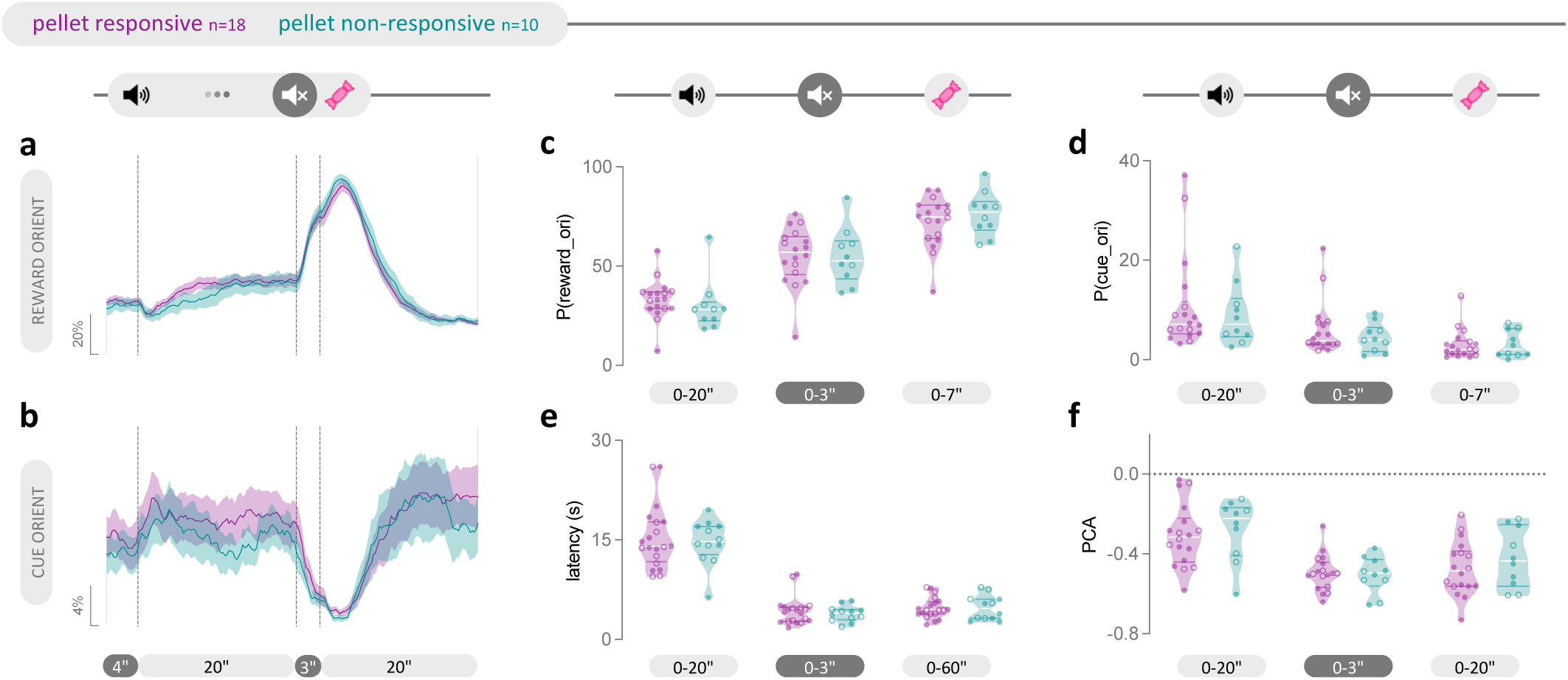
**Distinct salience encoding was not reflected in behavioral performance during probe testing. a-b**, Mean probability for reward-oriented (**a**) and cue-oriented (**b**) behavior in response to appetitive cue presentation and reward delivery during xRPE probe trials. **c-f**, No between-phenotype differences were observed in behavioral performance during probe sessions, including reward-oriented behavior (**c**), cue-oriented behavior (**d**), magazine entry latency (**e**), and PCA index (**f**).

**Supplemental Figure 7:**
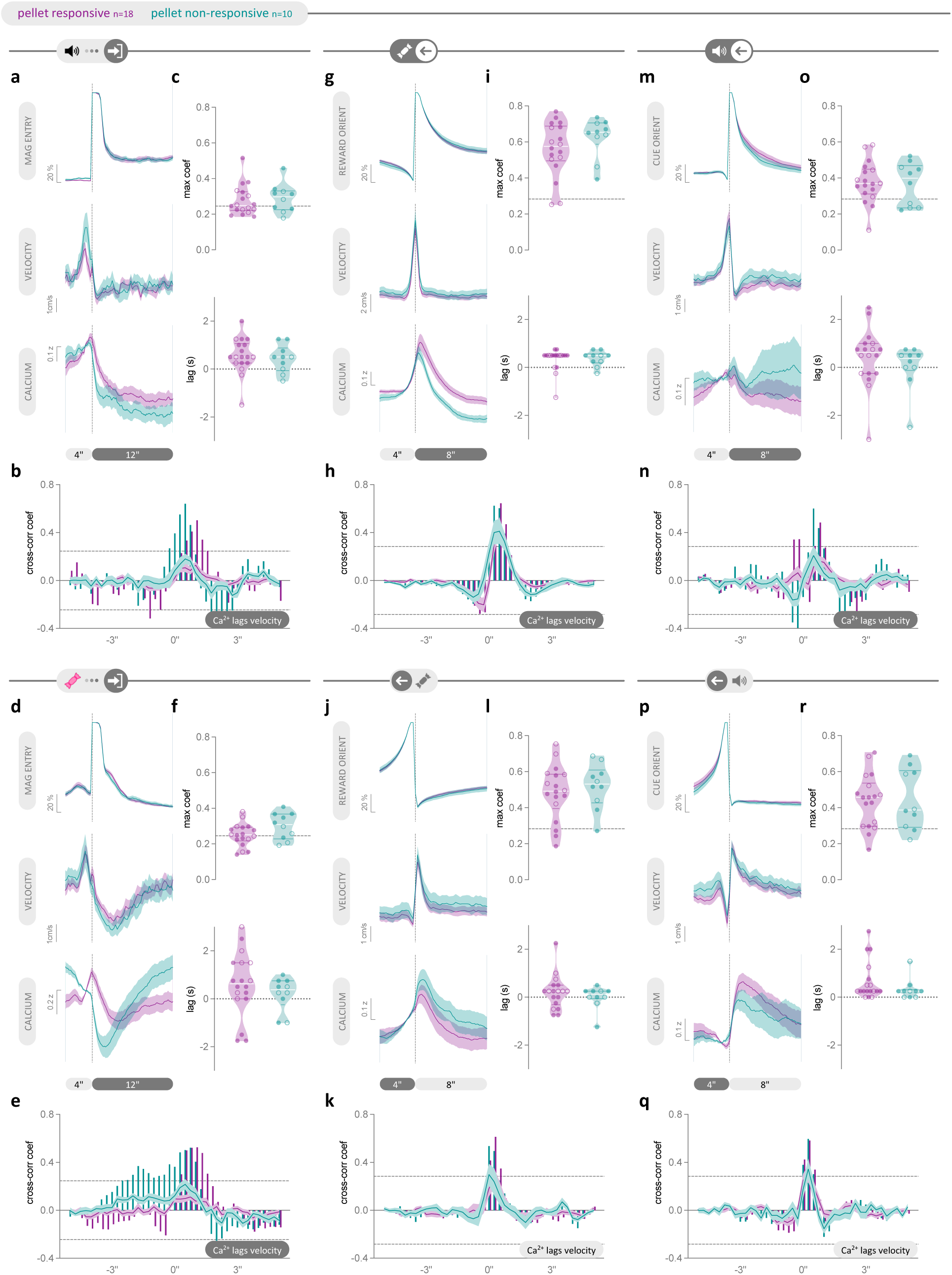
mPFC Ca^2+^ activity is consistently, positively correlated with movement across distinct appetitive behaviors and Ca^2+^ activity profiles. a, d, Mean magazine entry probability (top), velocity (middle), and mPFC Ca^2+^ activity (bottom) at the first magazine entry during cue presentation (a) and after pellet delivery (d) during xRPE probe trials. g, j, Mean probability of reward-oriented behavior (top), velocity (middle), and mPFC Ca^2+^ activity (bottom) at the onset (g) and offset (j) of entry to the food magazine zone during probe sessions. m, p, Mean probability of cue-oriented behavior (top), velocity (middle), and mPFC Ca^2+^ activity (bottom) at the onset (m) and offset (p) of entry into either cue zone during probe sessions. b, e, h, k, n, q, Velocity and mPFC Ca^2+^ activity are positively correlated with Ca^2+^ activity lagging all behaviors including the first magazine entry during cue presentation (b) and after pellet delivery (e), onset (h) and offset (k) of reward-oriented behavior, and onset (n) and offset (q) of cue-oriented behavior. c, f, i, l, o, r, The strength of the positive cross-correlation between velocity and mPFC Ca^2+^ (top) and the corresponding lag (bottom) did not differ between phenotypes for any measure. PSTHs are expressed as group mean ± shaded SEM. Violin plots indicate median and quartiles. Individual data are shown as circles (open = males, closed = females). Cross-correlograms show the cross-correlation function (XCF) between the group average velocity and mPFC Ca^2+^ activity (shown in bars), and the group average of individual rat’s XCF (shown in mean ± shaded SEM). Dashed lines on the cross-correlograms and violin plots indicate significance cutoff.

**Supplemental Figure 8.**
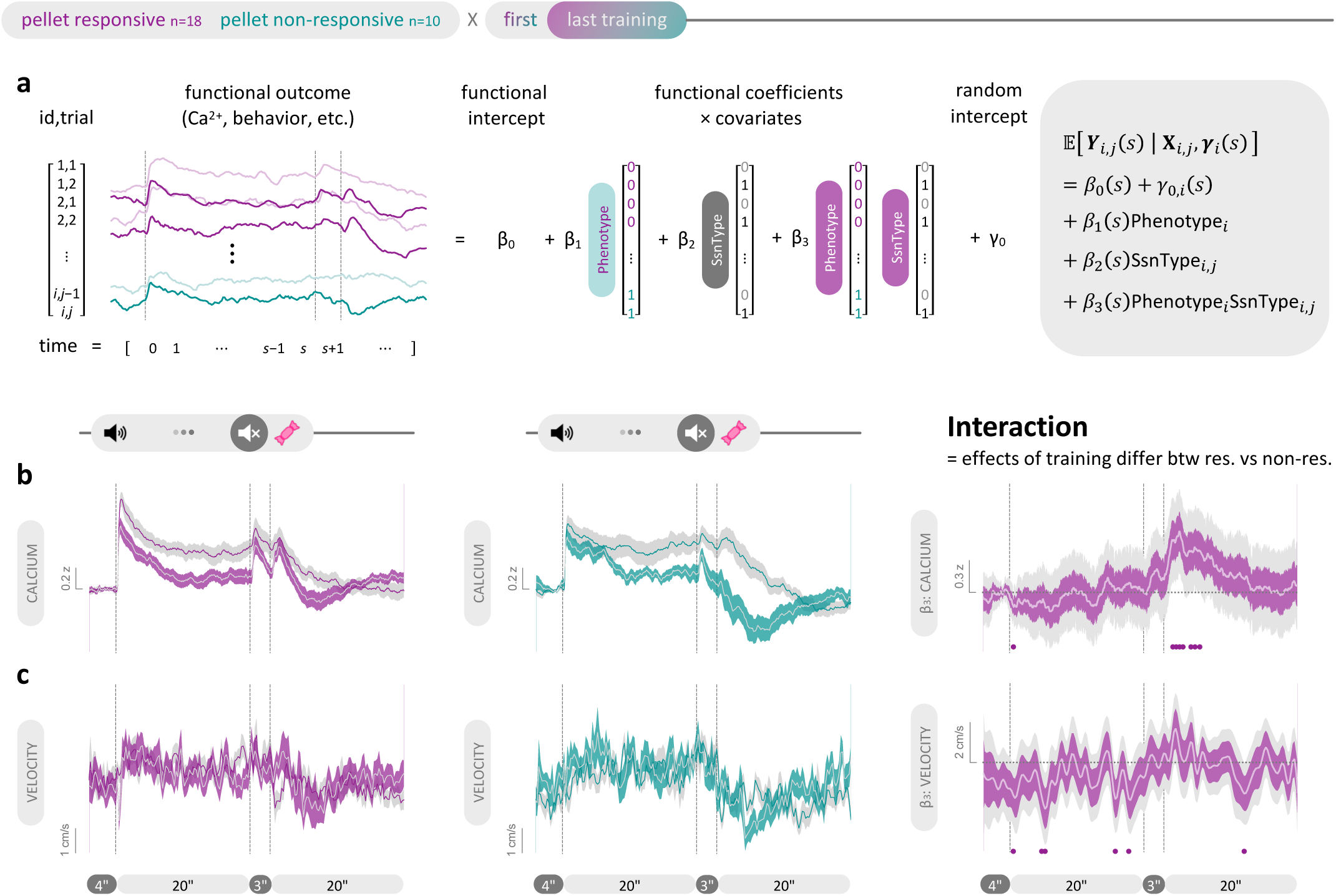
: Differences in pellet responsivity are not driven by differences in movement during training. **a**, Schematic (left) of FLMM (right) predicting various functional outcomes using session type and phenotype as covariates. **b**, Mean mPFC Ca^2+^ in response to cue presentation and pellet delivery during the first (grey shade) and last appetitive conditioning session (color shade) for the pellet-responsive (left) and non-responsive (middle) phenotypes. Right: Functional coefficient estimates showing that compared to the non-responsive phenotype, the responsive phenotype exhibited a significantly greater increase in mPFC Ca^2+^ in response to pellet delivery across training. **c**, Mean velocity in response to cue presentation and pellet delivery during the first (grey shade) and last appetitive conditioning session (color shade) for the pellet-responsive (left) and non-responsive (middle) phenotypes. Right: Functional coefficient estimates showing that compared to the non-responsive phenotype, pellet-responsive rats exhibited a larger decrease in movement during cue presentation and after pellet delivery across training. FLMM coefficients are expressed as estimates ± pointwise 95% CIs (dark shade) and joint 95% CIs (light shade) accounted for multiple comparisons. Statistical significance was accepted if the joint 95% CI of a timepoint did not contain y=0 (indicated by the horizontal dotted line) and was indicated by filled purple circles along the x axis.

**Supplemental Figure 9.**
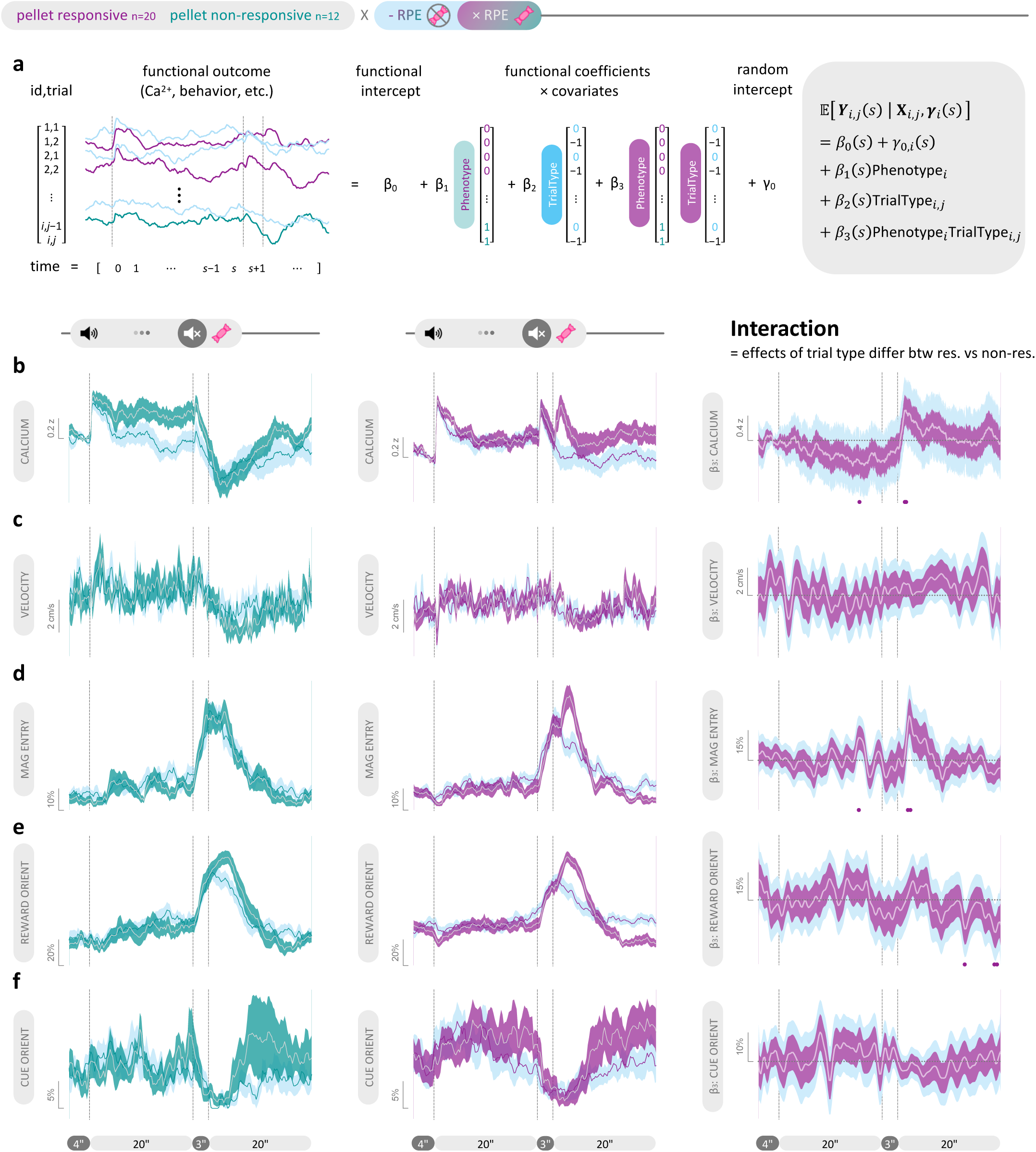
**: Distinct salience encoding supports adaptive reward seeking during reward omission. a**, Schematic (left) of FLMM (right) predicting various functional outcomes using trial type and phenotype as covariates. **b-f** Left and middle: Mean mPFC Ca^2+^ (**b**), velocity (**c**), magazine entry probability (**d**), reward-oriented behavior (**e**), and cue- oriented behavior (**f**) in response to cue presentation and pellet delivery during -RPE probe trials (cyan) and the subsequent xRPE trials (color) that followed for the pellet-responsive (left) and non-responsive (middle) phenotypes. **b,** Right: Functional coefficient estimates showing that compared to the non-responsive phenotype, the responsive phenotype exhibited a significantly larger increase in mPFC Ca^2+^ in response to pellet delivery relative to omission. **c**, Right: Functional coefficient estimates showing no difference in velocity between phenotypes as a result of trial type. **d**, Right: Functional coefficient estimates showing that the pellet-responsive phenotype exhibited a larger increase in reward seeking immediately after pellet delivery relative to omission compared to the non-responsive phenotype. **e**, Right: Functional coefficient estimates showing that the pellet-responsive phenotype exhibited a greater decrease in reward-oriented behavior during the ITI following pellet delivery relative to omission compared to the non-responsive phenotype. **f**, Right: Functional coefficient estimates showing no differences between phenotypes in cue-oriented behavior as a result of trial type. FLMM coefficients are expressed as estimates ± pointwise 95% CIs (dark shade) and joint 95% CIs (light shade) accounted for multiple comparisons. Statistical significance was accepted if the joint 95% CI of a timepoint did not contain y=0 (indicated by the horizontal dotted line) and was indicated by filled purple circles along the x axis.

**Supplemental Figure 10.**
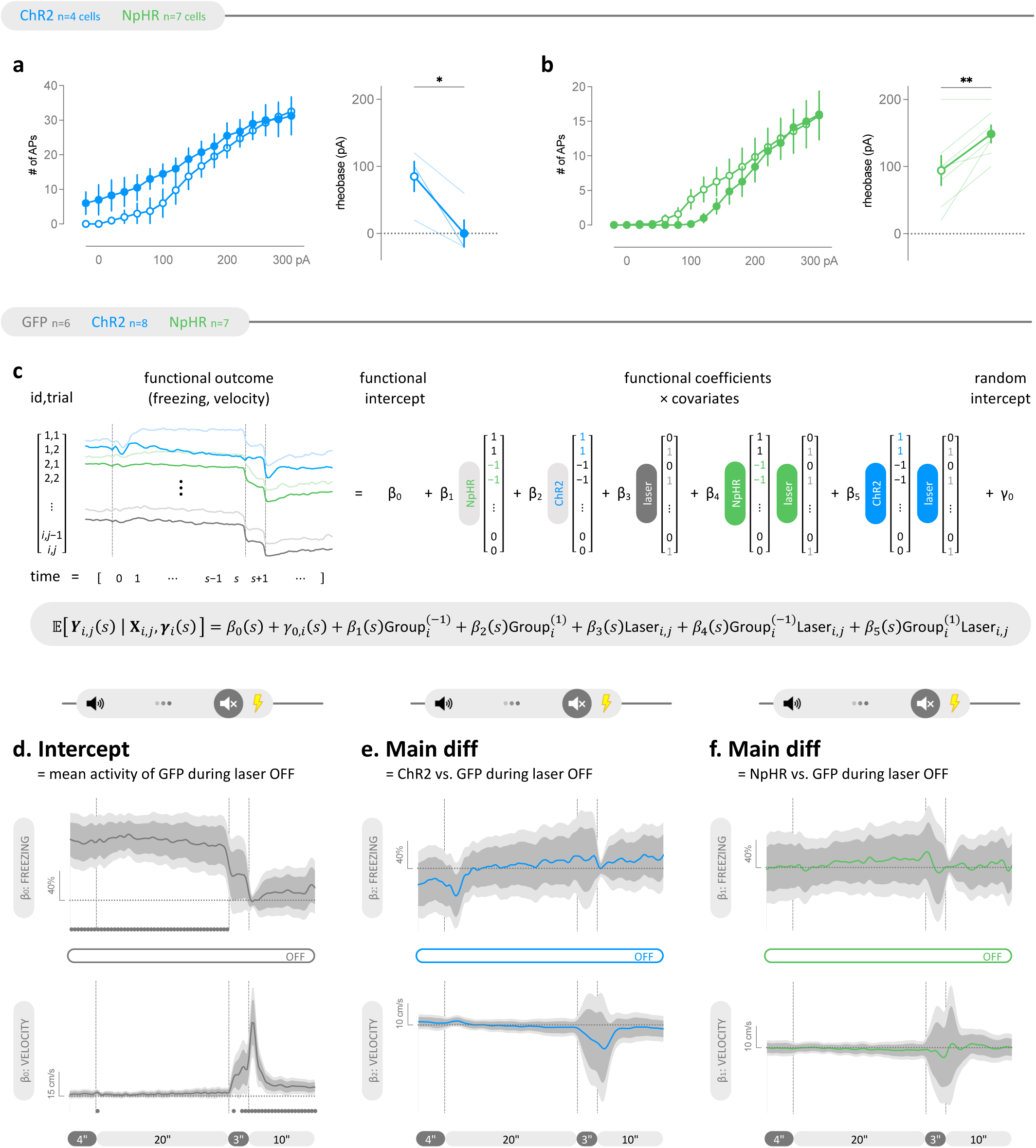
: Validation of optogenetic constructs and confirmation that experimental groups do not differ in baseline behavioral performance. a-b, Laser light delivery significantly reduced rheobase in ChR2-expressing mPFC neurons (a, paired t test, P=0.0259) and increased rheobase in NpHR-expressing mPFC neurons (b, paired t test, P=0.0090). c, Schematic (top) of FLMM (bottom) predicting freezing and movement using viral group and laser delivery as covariates. d, Functional intercept estimate showing significant freezing (top) before and during cue presentation, and significant movement (bottom) following cue offset and during shock presentation in GFP-expressing controls in the absence of laser light delivery. e-f, Functional coefficient estimate showing no significant difference in freezing (top) or movement (bottom) between ChR2-(e) or NpHR-expressing (f) rats vs. GFP-expressing controls in the absence of laser light delivery. Input-output curves are expressed as mean ± SEM (open symbols = laser OFF, closed symbols = laser ON). Before-after plots show both mean ± SEM and individual data. FLMM coefficients are expressed as estimates ± pointwise 95% CIs (dark shade) and joint 95% CIs (light shade) accounted for multiple comparisons. Statistical significance was accepted if the joint 95% CI of a timepoint did not contain y=0 (indicated by the horizontal dotted line) and was indicated by filled circles along the x axis. \**P* < 0.05, \*\**P* < 0.01.

**Supplemental Figure 11:**
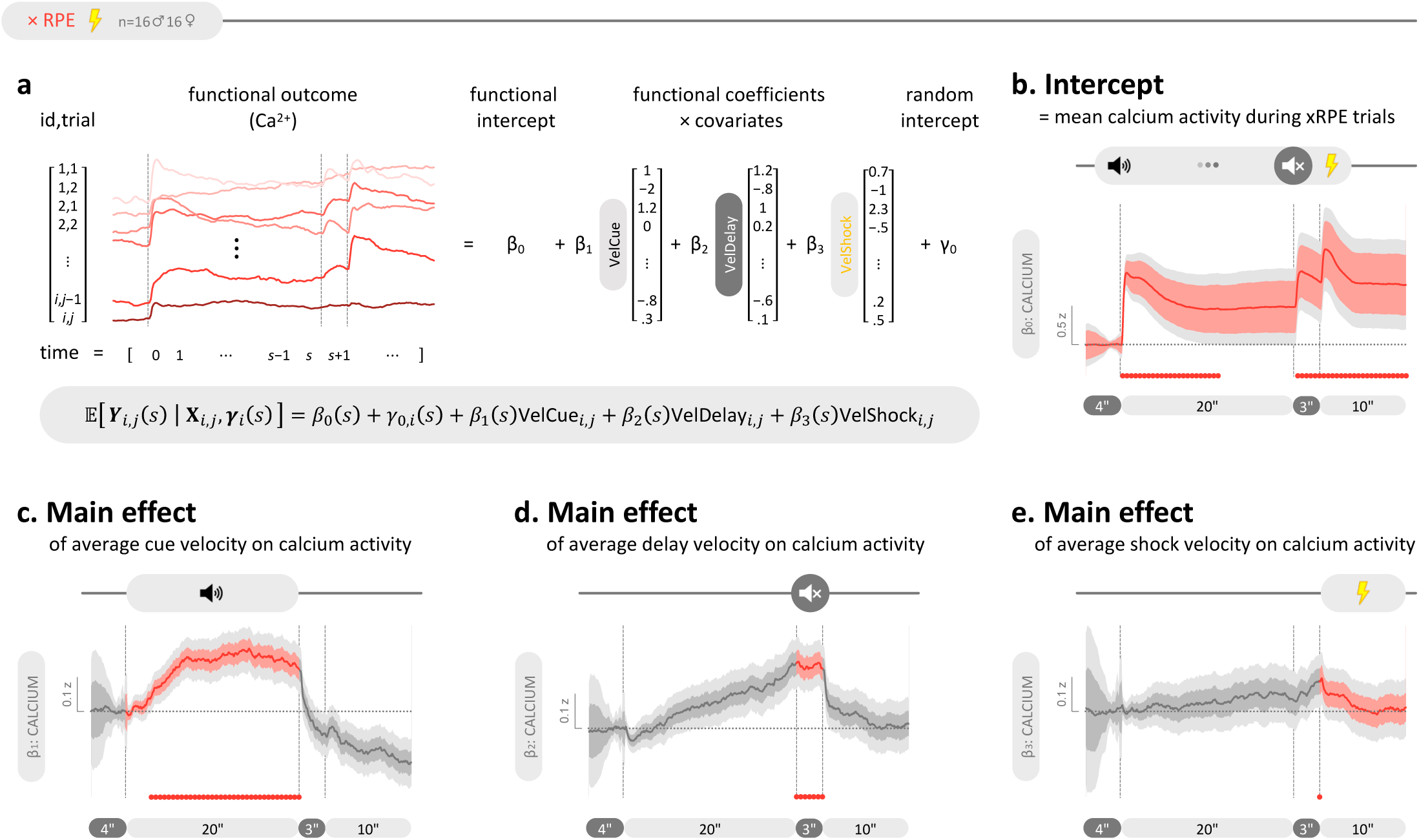
mPFC Ca^2+^ activity is significantly modulated by movement during shock-predicting cue presentation. **a**, Schematic (top) of FLMM (bottom) predicting mPFC Ca^2+^ activity using average velocity during each trial component as covariates to identify the movement-dominant periods for effective optogenetic manipulation. **b**, Functional intercept estimate confirms our empirical results showing increased mPFC Ca^2+^ activity during both cue presentation and shock delivery given average movement. **c-e**, Functional coefficient estimates showing a significant positive correlation between mPFC Ca^2+^ and velocity beginning 3 s after cue onset and lasting through cue offset at 20 s (**c**), as well as during the entire 3 s delay period (**d**), and the 0.5 s shock presentation period (**e**). FLMM coefficients are expressed as estimates ± pointwise 95% CIs (dark shade) and joint 95% CIs (light shade) accounted for multiple comparisons. Statistical significance was accepted if the joint 95% CI of a timepoint did not contain y=0 (indicated by the horizontal dotted line) and was indicated by filled red circles along the x axis.

## Supplemental

Functional bipartition of medial prefrontal cortex supports salience detection and movement gain

## Supplemental Information

**Supplemental Table 1:** Statistical analysis summary.

**Supplemental Movie 1:** Concurrent recording of mPFC Ca^2+^ activity and behavior.

**Supplemental Movie 2:** mPFC optogenetic stimulation drove generalized movement.

**Supplemental Movie 3:** Identification of freezing and darting using pose estimation.

**Supplemental Movie 4:** Identification of reward- and cue-orientation using pose estimation.

## References

1. Schultz, W., Dayan, P. & Montague, P. R. A neural substrate of prediction and reward. Science 275, 1593–1599 (1997).

2. Schultz, W. Dopamine reward prediction-error signalling: a two-component response. Nat Rev Neurosci 17, 183–195 (2016).

3. Redish, A. D. Addiction as a Computational Process Gone Awry. Science 306, 1944–1947 (2004).

4. Whitton, A. E., Treadway, M. T. & Pizzagalli, D. A. Reward processing dysfunction in major depression, bipolar disorder and schizophrenia. Curr Opin Psychiatry 28, 7–12 (2015).

5. Yaple, Z. A., Tolomeo, S. & Yu, R. Abnormal prediction error processing in schizophrenia and depression. Hum Brain Mapp 42, 3547–3560 (2021).

6. Millard, S. J., Bearden, C. E., Karlsgodt, K. H. & Sharpe, M. J. The prediction-error hypothesis of schizophrenia: new data point to circuit-specific changes in dopamine activity. Neuropsychopharmacology 47, 628–640 (2022).

7. Yang, X., Song, Y., Zou, Y., Li, Y. & Zeng, J. Neural correlates of prediction error in patients with schizophrenia: evidence from an fMRI meta-analysis. Cereb Cortex 34, bhad471 (2024).

8. Rutledge, R. B. et al. Association of Neural and Emotional Impacts of Reward Prediction Errors With Major Depression. JAMA Psychiatry 74, 790–797 (2017).

9. Kumar, P. et al. Impaired reward prediction error encoding and striatal-midbrain connectivity in depression. Neuropsychopharmacology 43, 1581–1588 (2018).

10. Norman, L. J. et al. Error Processing and Inhibitory Control in Obsessive-Compulsive Disorder: A Meta-analysis Using Statistical Parametric Maps. Biol Psychiatry 85, 713–725 (2019).

11. Parvaz, M. A. et al. Impaired neural response to negative prediction errors in cocaine addiction. J. Neurosci. 35, 1872–1879 (2015).

12. Tolomeo, S., Yaple, Z. A. & Yu, R. Neural representation of prediction error signals in substance users. Addict Biol 26, e12976 (2021).

13. Konova, A. B. et al. Reduced neural encoding of utility prediction errors in cocaine addiction. Neuron 111, 4058–4070.e6 (2023).

14. Costello, H. et al. Disrupted reward processing in Parkinson’s disease and its relationship with dopamine state and neuropsychiatric syndromes: a systematic review and meta-analysis. J Neurol Neurosurg Psychiatry 93, 555–562 (2022).

15. Mosner, M. G. et al. Neural Mechanisms of Reward Prediction Error in Autism Spectrum Disorder. Autism Res Treat 2019, 5469191 (2019).

16. Garrison, J., Erdeniz, B. & Done, J. Prediction error in reinforcement learning: a meta-analysis of neuroimaging studies. Neurosci Biobehav Rev 37, 1297–1310 (2013).

17. Chase, H. W., Kumar, P., Eickhoff, S. B. & Dombrovski, A. Y. Reinforcement learning models and their neural correlates: An activation likelihood estimation meta-analysis. Cogn Affect Behav Neurosci 15, 435–459 (2015).

18. Corlett, P. R., Mollick, J. A. & Kober, H. Meta-analysis of human prediction error for incentives, perception, cognition, and action. Neuropsychopharmacology 47, 1339–1349 (2022).

19. Rushworth, M. F. S. & Behrens, T. E. J. Choice, uncertainty and value in prefrontal and cingulate cortex. Nat Neurosci 11, 389–397 (2008).

20. Alexander, W. H. & Brown, J. W. Medial prefrontal cortex as an action-outcome predictor. Nat Neurosci 14, 1338–1344 (2011).

21. Alexander, W. H. & Brown, J. W. The Role of the Anterior Cingulate Cortex in Prediction Error and Signaling Surprise. Top Cogn Sci 11, 119–135 (2019).

22. Kennerley, S. W., Behrens, T. E. J. & Wallis, J. D. Double dissociation of value computations in orbitofrontal and anterior cingulate neurons. Nat Neurosci 14, 1581–1589 (2011).

23. Hyman, J. M., Holroyd, C. B. & Seamans, J. K. A Novel Neural Prediction Error Found in Anterior Cingulate Cortex Ensembles. Neuron 95, 447–456.e3 (2017).

24. Fu, Z., Sajad, A., Errington, S. P., Schall, J. D. & Rutishauser, U. Neurophysiological mechanisms of error monitoring in human and non-human primates. Nat Rev Neurosci 1–20 (2023) doi:10.1038/s41583-022-00670-w.

25. Hoy, C. W. et al. Asymmetric coding of reward prediction errors in human insula and dorsomedial prefrontal cortex. Nat Commun 14, 8520 (2023).

26. Cole, N., Harvey, M., Myers-Joseph, D., Gilra, A. & Khan, A. G. Prediction-error signals in anterior cingulate cortex drive task-switching. Nat Commun 15, 7088 (2024).

27. Der-Avakian, A. et al. Identification of conserved frontal neurophysiological markers of cognitive flexibility in humans and rats. Commun Biol 8, 1268 (2025).

28. Holroyd, C. B. & Coles, M. G. H. The neural basis of human error processing: Reinforcement learning, dopamine, and the error-related negativity. Psychological Review 109, 679–709 (2002).

29. Hughes, N. C. Reward circuit local field potential modulations precede risk taking.

30. Oya, H. et al. Electrophysiological correlates of reward prediction error recorded in the human prefrontal cortex. Proc. Natl. Acad. Sci. U.S.A. 102, 8351–8356 (2005).

31. Miller, E. K. The prefrontal cortex and cognitive control. Nat Rev Neurosci 1, 59–65 (2000).

32. Howland, J. G., Ito, R., Lapish, C. C. & Villaruel, F. R. The rodent medial prefrontal cortex and associated circuits in orchestrating adaptive behavior under variable demands. Neurosci Biobehav Rev 135, 104569 (2022).

33. Ji, Q. et al. Role of prefrontal cortex in freezing of gait in Parkinson’s disease: mechanisms and neuroimaging evidence. Front Neurol 17, 1713795 (2026).

34. Mediane, D. H., Basu, S., Cahill, E. N. & Anastasiades, P. G. Medial prefrontal cortex circuitry and social behaviour in autism. Neuropharmacology 260, 110101 (2024).

35. Van den Oever, M. C., Spijker, S., Smit, A. B. & De Vries, T. J. Prefrontal cortex plasticity mechanisms in drug seeking and relapse. Neurosci Biobehav Rev 35, 276–284 (2010).

36. Elliott, R., Zahn, R., Deakin, J. F. W. & Anderson, I. M. Affective cognition and its disruption in mood disorders. Neuropsychopharmacology 36, 153–182 (2011).

37. Sherathiya, V. N., Schaid, M. D., Seiler, J. L., Lopez, G. C. & Lerner, T. N. GuPPy, a Python toolbox for the analysis of fiber photometry data. Sci Rep 11, 24212 (2021).

38. Kriegeskorte, N., Mur, M. & Bandettini, P. A. Representational similarity analysis - connecting the branches of systems neuroscience. Front. Syst. Neurosci. 2, (2008).

39. Liu, Y. et al. The Mesolimbic Dopamine Activity Signatures of Relapse to Alcohol-Seeking. J. Neurosci. 40, 6409–6427 (2020).

40. Mahn, M., Prigge, M., Ron, S., Levy, R. & Yizhar, O. Biophysical constraints of optogenetic inhibition at presynaptic terminals. Nat Neurosci 19, 554–556 (2016).

41. Hou, S. & Glover, E. J. Pi USB Cam: A Simple and Affordable DIY Solution That Enables High-Quality, High-Throughput Video Capture for Behavioral Neuroscience Research. eNeuro 9, (2022).

42. Pereira, T. D. et al. SLEAP: A deep learning system for multi-animal pose tracking. Nat Methods 19, 486–495 (2022).

43. Gabriel, C. J. et al. BehaviorDEPOT is a simple, flexible tool for automated behavioral detection based on markerless pose tracking. eLife 11, e74314 (2022).

44. Bohnslav, J. P. et al. DeepEthogram, a machine learning pipeline for supervised behavior classification from raw pixels. eLife 10, e63377 (2021).

45. Gruene, T. M., Flick, K., Stefano, A., Shea, S. D. & Shansky, R. M. Sexually divergent expression of active and passive conditioned fear responses in rats. eLife 4, e11352 (2015).

46. Flagel, S. B. et al. A food predictive cue must be attributed with incentive salience for it to induce c-fos mRNA expression in cortico-striatal-thalamic brain regions. Neuroscience 196, 80–96 (2011).

47. Meyer, P. J. et al. Quantifying Individual Variation in the Propensity to Attribute Incentive Salience to Reward Cues. PLOS ONE 7, e38987 (2012).

48. Iglesias, A. G. et al. Inhibition of Dopamine Neurons Prevents Incentive Value Encoding of a Reward Cue: With Revelations from Deep Phenotyping. J. Neurosci. 43, 7376–7392 (2023).

49. Loewinger, G., Cui, E., Lovinger, D. & Pereira, F. A statistical framework for analysis of trial-level temporal dynamics in fiber photometry experiments. eLife 13, RP95802 (2025).

50. Musall, S., Kaufman, M. T., Juavinett, A. L., Gluf, S. & Churchland, A. K. Single-trial neural dynamics are dominated by richly varied movements. Nat Neurosci 22, 1677–1686 (2019).

51. Seeley, W. W. et al. Dissociable Intrinsic Connectivity Networks for Salience Processing and Executive Control. J. Neurosci. 27, 2349–2356 (2007).

52. Uddin, L. Q. Salience processing and insular cortical function and dysfunction. Nat Rev Neurosci 16, 55–61 (2015).

53. Kalhan, S., Redish, A. D., Hester, R. & Garrido, M. I. A salience misattribution model for addictive-like behaviors. Neuroscience & Biobehavioral Reviews 125, 466–477 (2021).

54. Winke, N., Lüthi, A., Herry, C. & Jercog, D. Prefrontal neural geometry of learned cues guides motivated behaviours. Nature 1–10 (2026) doi:10.1038/s41586-025-09902-2.

55. Zhang, K. et al. Prefrontal chandelier cells encode stimulus salience to influence learning in male mice. Nat Commun 10.1038/s41467-026-68959-3 (2026) doi:10.1038/s41467-026-68959-3.

56. Peters, J., Kalivas, P. W. & Quirk, G. J. Extinction circuits for fear and addiction overlap in prefrontal cortex. Learn. Mem. 16, 279–288 (2009).

57. Bravo-Rivera, C., Roman-Ortiz, C., Brignoni-Perez, E., Sotres-Bayon, F. & Quirk, G. J. Neural Structures Mediating Expression and Extinction of Platform-Mediated Avoidance. J. Neurosci. 34, 9736–9742 (2014).

58. Courtin, J. et al. Prefrontal parvalbumin interneurons shape neuronal activity to drive fear expression. Nature 505, 92–96 (2014).

59. Dejean, C. et al. Prefrontal neuronal assemblies temporally control fear behaviour. Nature 535, 420–424 (2016).

60. Gourley, S. L. & Taylor, J. R. Going and stopping: dichotomies in behavioral control by the prefrontal cortex. Nat Neurosci 19, 656–664 (2016).

61. Karalis, N. et al. 4-Hz oscillations synchronize prefrontal–amygdala circuits during fear behavior. Nat Neurosci 19, 605–612 (2016).

62. Burgos-Robles, A. et al. Amygdala inputs to prefrontal cortex guide behavior amid conflicting cues of reward and punishment. Nat Neurosci 20, 824–835 (2017).

63. Otis, J. M. et al. Prefrontal cortex output circuits guide reward seeking through divergent cue encoding. Nature 543, 103–107 (2017).

64. Diehl, M. M. et al. Active avoidance requires inhibitory signaling in the rodent prelimbic prefrontal cortex. eLife 7, e34657 (2018).

65. Vander Weele, C. M. et al. Dopamine enhances signal-to-noise ratio in cortical-brainstem encoding of aversive stimuli. Nature 563, 397–401 (2018).

66. Bagur, S. et al. Breathing-driven prefrontal oscillations regulate maintenance of conditioned-fear evoked freezing independently of initiation. Nat Commun 12, 2605 (2021).

67. Jercog, D. et al. Dynamical prefrontal population coding during defensive behaviours. Nature 595, 690–694 (2021).

68. Kajs, B. L., Loewke, A. C., Dorsch, J. M., Vinson, L. T. & Gunaydin, L. A. Divergent encoding of active avoidance behavior in corticostriatal and corticolimbic projections. Sci Rep 12, 10731 (2022).

69. Ehret, B. et al. Population-level coding of avoidance learning in medial prefrontal cortex. Nat Neurosci 27, 1805–1815 (2024).

70. Dana, H. et al. High-performance calcium sensors for imaging activity in neuronal populations and microcompartments. Nat Methods 16, 649–657 (2019).

71. Stringer, C. et al. Spontaneous Behaviors Drive Multidimensional, Brain-wide Activity. Science 364, 255 (2019).

72. Yager, L. M., Pitchers, K. K., Flagel, S. B. & Robinson, T. E. Individual Variation in the Motivational and Neurobiological Effects of an Opioid Cue. Neuropsychopharmacol 40, 1269–1277 (2015).

73. Colaizzi, J. M. et al. Mapping sign-tracking and goal-tracking onto human behaviors. Neurosci Biobehav Rev 111, 84–94 (2020).

74. Colaizzi, J. M. et al. The propensity to sign-track is associated with externalizing behavior and distinct patterns of reward-related brain activation in youth. Sci Rep 13, 4402 (2023).

75. Robinson, T. E. & Flagel, S. B. Dissociating the Predictive and Incentive Motivational Properties of Reward-Related Cues Through the Study of Individual Differences. Biological Psychiatry 65, 869–873 (2009).

76. Saunders, B. T. & Robinson, T. E. A Cocaine Cue Acts as an Incentive Stimulus in Some but not Others: Implications for Addiction. Biological Psychiatry 67, 730–736 (2010).

77. Robinson, T. E., Yager, L. M., Cogan, E. S. & Saunders, B. T. On the motivational properties of reward cues: Individual differences. Neuropharmacology 76, 450–459 (2014).

78. Steinmetz, N. A., Zatka-Haas, P., Carandini, M. & Harris, K. D. Distributed coding of choice, action and engagement across the mouse brain. Nature 576, 266–273 (2019).

79. International Brain Laboratory et al. A brain-wide map of neural activity during complex behaviour. Nature 645, 177–191 (2025).

80. Wang, Z. A. et al. Brain-wide analysis reveals movement encoding structured across and within brain areas. Nat Neurosci 29, 147–158 (2026).

81. Fadok, J. P. et al. A competitive inhibitory circuit for selection of active and passive fear responses. Nature 542, 96–100 (2017).

82. Martin-Fernandez, M. et al. Prefrontal circuits encode both general danger and specific threat representations. Nat Neurosci 26, 2147–2157 (2023).

83. Weinreb, C. et al. Spontaneous behavior is a succession of self-directed tasks. Neuron 114, 922–937.e12 (2026).

84. Kutlu, M. G. et al. Dopamine release in the nucleus accumbens core signals perceived saliency. Current Biology 31, 4748–4761.e8 (2021).

85. Bakhurin, K. et al. Dopamine dynamics during stimulus-reward learning in mice can be explained by performance rather than learning. Nat Commun 16, 9081 (2025).

86. Jacobs, D. S. & Moghaddam, B. Prefrontal Cortex Representation of Learning of Punishment Probability During Reward-Motivated Actions. J. Neurosci. 40, 5063–5077 (2020).

87. Simpson, E. H. et al. Lights, fiber, action! A primer on in vivo fiber photometry. Neuron 112, 718–739 (2024).

88. Hua, S.-S. et al. NMDA Receptor–Dependent Synaptic Potentiation via APPL1 Signaling Is Required for the Accessibility of a Prefrontal Neuronal Assembly in Retrieving Fear Extinction. Biological Psychiatry 94, 262–277 (2023).

89. Jacobs, D. S., Allen, M. C., Park, J. & Moghaddam, B. Learning of probabilistic punishment as a model of anxiety produces changes in action but not punisher encoding in the dmPFC and VTA. eLife 11, e78912 (2022).

90. Liu, X.-D. et al. Hippocampal Lnx1–NMDAR multiprotein complex mediates initial social memory. Mol Psychiatry 26, 3956–3969 (2021).

91. Sias, A. C. et al. A bidirectional corticoamygdala circuit for the encoding and retrieval of detailed reward memories. eLife 10, e68617 (2021).

92. Mante, V., Sussillo, D., Shenoy, K. V. & Newsome, W. T. Context-dependent computation by recurrent dynamics in prefrontal cortex. Nature 503, 78–84 (2013).

93. Fusi, S., Miller, E. K. & Rigotti, M. Why neurons mix: high dimensionality for higher cognition. Current Opinion in Neurobiology 37, 66–74 (2016).

94. Tye, K. M. et al. Mixed selectivity: Cellular computations for complexity. Neuron 112, 2289– 2303 (2024).

95. Tafazoli, S. et al. Building compositional tasks with shared neural subspaces. Nature 650, 164–172 (2026).

96. Hasnain, M. A. et al. Separating cognitive and motor processes in the behaving mouse. Nat Neurosci 28, 640–653 (2025).

97. Gallello, I. et al. Dopamine responses in medial frontal cortex are more consistent with a generalized arousal signal than signed reward prediction errors. J. Neurosci. (2026) doi:10.1523/JNEUROSCI.1454-25.2026.

98. Rigotti, M. et al. The importance of mixed selectivity in complex cognitive tasks. Nature 497, 585–590 (2013).

99. Xie, Y. et al. Geometry of sequence working memory in macaque prefrontal cortex. Science 375, 632–639 (2022).

100. Jazayeri, M. & Afraz, A. Navigating the Neural Space in Search of the Neural Code. Neuron 93, 1003–1014 (2017).

101. Wolff, S. B. E. & Ölveczky, B. P. The promise and perils of causal circuit manipulations. Curr Opin Neurobiol 49, 84–94 (2018).

102. Bertrand, O. C. et al. Brawn before brains in placental mammals after the end-Cretaceous extinction. Science 376, 80–85 (2022).

103. Schwartz, E. et al. Evolution of cortical geometry and its link to function, behaviour and ecology. Nat Commun 14, 2252 (2023).

104. Bertrand, O. C. & Krubitzer, L. The functional adaptations of mammalian brain structures through a behavioural ecology lens. Nat. Rev. Biodivers. 1, 703–716 (2025).

105. Smaers, J. B. et al. The evolution of mammalian brain size. Science Advances 7, eabe2101 (2021).

106. Manley, J. et al. Simultaneous, cortex-wide dynamics of up to 1 million neurons reveal unbounded scaling of dimensionality with neuron number. Neuron 112, 1694–1709.e5 (2024).

